# Synthetic α-synuclein fibrils replicate in mice causing MSA-like pathology

**DOI:** 10.1101/2024.07.01.601498

**Authors:** Domenic Burger, Marianna Kashyrina, Lukas van den Heuvel, Hortense de La Seiglière, Amanda J. Lewis, Francesco De Nuccio, Inayathulla Mohammed, Jérémy Verchère, Cécile Feuillie, Mélanie Berbon, Marie-Laure Arotcarena, Aude Retailleau, Erwan Bezard, Marie-Hélène Canron, Wassilios G. Meissner, Antoine Loquet, Luc Bousset, Christel Poujol, K. Peter R. Nilsson, Florent Laferrière, Thierry Baron, Dario Domenico Lofrumento, Francesca De Giorgi, Henning Stahlberg, François Ichas

## Abstract

Multiple system atrophy (MSA) is a rapidly progressive neurodegenerative disease of unknown cause, typically affecting individuals aged 50–60 and leading to death within a decade^1–3^. It is characterized by glial cytoplasmic inclusions (GCIs) composed of fibrillar alpha-synuclein (aSyn)^4–8^, whose formation shows parallels with prion propagation^9,10^. While fibrils extracted from MSA brains have been structurally characterized^11^, their ability to replicate in a “protein-only” manner has been questioned^12^, and their capacity to induce GCIs *in vivo* remains unexplored. By contrast, the synthetic fibril strain 1B^13,14^, assembled from recombinant human aSyn, self-replicates *in vitro* and induces GCIs in mice^15^ - suggesting direct relevance to MSA - but awaited scrutiny at an atomic scale. Here, we report high-resolution structural analyses of 1B fibrils and of fibrils extracted from diseased mice injected with 1B that developed GCIs (1B^P^). We show *in vivo* that conformational templating enables fibril strain replication, resulting in MSA-like inclusion pathology. Remarkably, the structures of 1B and 1B^P^ are highly similar and mimic the fold of aSyn observed in one protofilament of fibrils isolated from MSA patients^11^. Moreover, reinjection of crude mouse brain homogenates containing 1B^P^ into new mice reproduces the same MSA-like pathology induced by the parent synthetic seed 1B. Our findings identify 1B as a synthetic pathogen capable of self-replication *in vivo* and reveal structural features of 1B/1B^P^ that may underlie MSA pathology, offering insights for therapeutic strategies.

## Body Text

MSA is characterized by the rapid invasion of the brainstem, the cerebellum and/or the basal ganglia by intracellular inclusions filled with fibrillar aSyn^4–8^. These inclusions appear in neurons and in oligodendrocytes (OLs), particularly within the myelinated bundles of the white matter^16–18^. The latter inclusions are called GCIs and are specific to MSA. They are only observed rarely in other sporadic alpha-synucleinopathies^1,2^. The phenomenon of a “step-by-step” progression that characterizes brain invasion by GCIs, and the speed at which the GCIs induce the death of neurons in myelinated tracts^1^, has led to the notion that MSA may obey a molecular mechanism analogous to that governing the intracerebral diffusion of pathogenic prions^9,10^. According to this hypothesis, the specific aSyn fibril strain found in GCIs plays a central role as both initiator and propagator of the pathology^9,10^.

aSyn fibrils in GCIs were first observed *in situ*^4–8^, and extracted and described using electron microscopy more than 25 years ago^19,20^. Such fibrils, extracted using the detergent sarkosyl, were characterized at high resolution by cryo-transmission electron microscopy (cryo-EM)^11^, but to date, it has not been demonstrated that they can self-replicate like prions or induce the formation of GCIs in animal models. The hypothesis was put forward^12^ that unknown cofactors might be required for the replication of these extracted assemblies.

In parallel to the extractive approach^11^, three synthetic aSyn fibril strains were obtained from recombinant aSyn, and these fibrils could self-replicate *in vitro* without the need for an external cofactor, and induce GCIs in mice. Chronologically, the first strain was a non-helical double-filament fibrillar assembly (ribbons) composed of monomers of human aSyn, capable of inducing GCIs, but only in rats overexpressing human aSyn concomitantly^21^. The second strain corresponded to mouse synthetic aSyn fibrils, which could induce the delayed appearance of GCIs in non-transgenic mice^22^. Finally, the third case concerned a fibril strain that some of us isolated and characterized (strain 1B), formed of human recombinant aSyn and causing the rapid appearance of GCIs in interfascicular OLs of wild-type mice^15^. Unlike for the aSyn fibrils extracted from MSA patients, no high-resolution structure of these types of synthetic aSyn fibrils had so far been available. This prevented the comparison of these fibrils with the atomic models obtained from patients to understand the structural bases of their capacity to induce GCIs.

We here report the structural characterization of aSyn fibrils of strain 1B at a resolution of 1.94 Å, and compare them to the aSyn filaments extracted from the brains of 1B-injected diseased mice in which GCIs formed (1B^P^, resolved at 3.8 Å). The striking structural similitude between 1B and 1B^P^ demonstrates that a conformational templating and replication process is indeed taking place *in vivo* and that it is responsible for fibril growth and inclusion pathology buildup. Further, the structural analogy of the 1B and 1B^P^ fold with the human MSA fold found in patient 5 of the seminal study of Goedert’s group^11^ suggests that a similar conformational replication process underlies MSA in humans.

## MSA-like neuropathology seeded by 1B

1B fibrils present a typical rod-like compact appearance with a width compatible with the intertwining of two protofilaments^13^ and a left-handed helicoidal twist (**Extended data Fig. 1a)**. We observed in our founder experiments that 1B was capable of seeding a particularly strong aSyn pathology in neuronal cultures (see also **Extended data Fig. 1b**) and *in vivo* in mice^13,14^. In this early work, we found that the pathology induced by 1B at early time points was made of Lewy neurites, neuronal cytoplasmic inclusions (NCIs), and most strikingly, of neuronal intranuclear inclusions (NIIs)^13,14^. Together with GCIs and Glial Nuclear Inclusions (GNIs), NIIs constitute a key feature of MSA pathology, i.e., they are distinctive for MSA compared to Parkinson’s disease or dementia with Lewy bodies (DLB), in which NIIs are absent^23,24^. This very cytopathological feature led us to investigate the possibility that the synthetic 1B fibrils could represent an MSA-like fibril strain. Using a directed anatomical approach and later time points, we observed that 6 months after intracerebral inoculation, 1B was indeed capable of also causing the appearance of GCIs in the anterior commissure (AC) of wild-type mice^15^, *i.e.*, in a major myelinated tract containing densely packed rows of interfascicular OLs. The time course and the prominence of the GCI pathology seeded in this tract supported the notion that like in MSA^24,25^, the aSyn pathology seeded by 1B fibrils originated in neurons and was secondarily propagated to OLs^15^.

After 24 months, a proper “invasion” of the posterior limb of the AC and of the bed nucleus of the *stria terminalis* (BNST) by GCIs could be observed (**Fig. 1a-e**), while Iba1-positive microglia did not particularly cluster around aSyn inclusion pathology (**Fig. 1c,d**). Confocal 3D volume reconstructions of affected OLs in the BNST showed the typical “flame-like” cytoplasmic distribution of phosphorylated aSyn (**Fig. 1e**). Global distribution of GCIs in full horizontal brain sections from animals sacrificed 1.5, 12, or 24 months after 1B fibril injection in their right caudate putamen (CP) showed generalized spread of GCIs (**Fig. 1f**, green markers). This was particularly prominent in the animal sacrificed at 24 months: in addition to the AC and to the BNST, GCIs were found bilaterally in deep cortical layers and the subcortical white matter, the CP, and unilaterally in the *globus pallidus* (GP) (injection side), the internal capsule (injection side), the entopeduncular nucleus (injection side), the *substantia nigra* (SN) *pars reticulata* (injection side), and the medullary reticular nucleus (contralateral side). A hovering section overview corresponding to **Fig. 1a-e** can be found in the **Supplementary Information Video-1**.

**Fig. 1.**
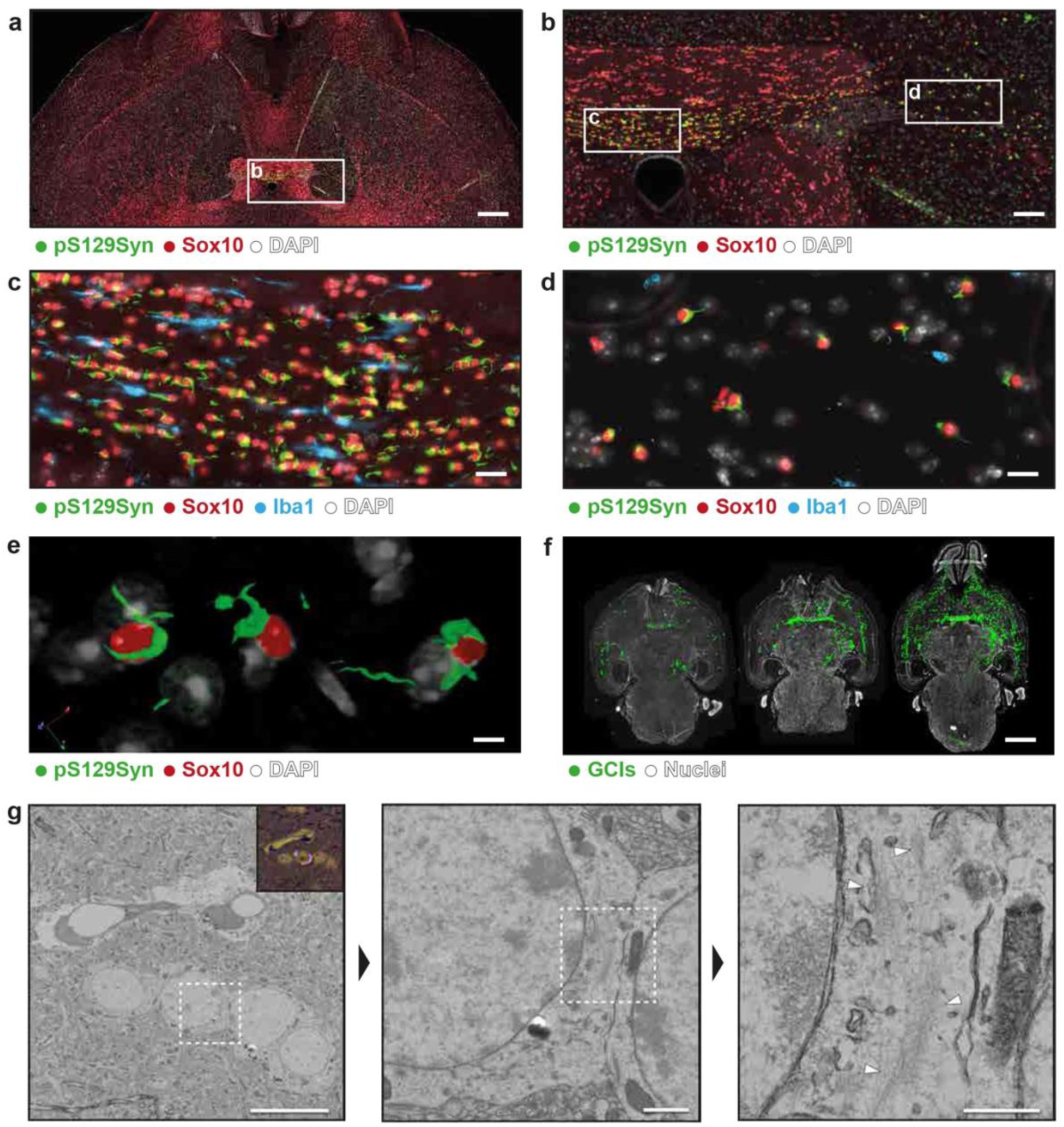
Seeding of GCIs and fibrillar pathology in wild-type mice by alpha-synuclein fibrils of the 1B strain. **a.** Topographic immunofluorescence of a horizontal brain section from a wild-type mouse injected 24 months earlier in the right CP with 1B fibrils. aSyn inclusions are labeled with anti-phospho aSyn (#64, green), OLs with anti-Sox10 (red), and nuclei with DAPI (white). The section intercepts the AC and BNST (box). **b.** Higher magnification of the boxed region showing invasion of the posterior limb of the AC (c) and BNST (d) by aSyn inclusions. **c.** Posterior limb of the AC: most interfascicular OLs harbor aSyn inclusions, i.e., GCIs. Sparse Iba1-positive microglia (blue) are also visible. **d.** BNST: Sox10-positive OLs contain aSyn inclusions (GCIs); sparse Iba1-positive microglia are detectable. **e.** Confocal 3D reconstruction centered on three OLs of the BNST bearing GCIs, from the section shown in (a). See also Supplementary Information Video-1. **f.** Neuroanatomical distribution of GCIs in brains at 1.5 (left), 12 (center), and 24 months (right) post-1B injection into the right CP. Whole-brain horizontal sections with every GCI manually annotated based on pS129Syn and Sox10. The 24-month section corresponds to the n+1 section adjacent to (a). **g.** Correlative light and electron microscopy (LM, EM) of pS129Syn-positive regions. From left to right: EM overview with LM correlation inset; closer view of pathology; and high magnification used for tomography to confirm fibrillar material. White arrowheads indicate fibrils. LM inset shows the same region as EM with IHC/toluidine blue overlay and inverted color for clarity. Scale bars: **a** = 500 µm; **b** = 100 µm; **c,d** = 20 µm; **e** = 5 µm; **f** = 3 mm; **g** (left–right) = 10 µm, 1 µm, 0.5 µm.

Depending on the neuroanatomical structure and the time point under consideration, the experimental GCIs of **Fig. 1f** coexisted with a variable load of surrounding neuronal somatic aSyn inclusions (NCIs and NIIs), which represent the forefront of the somatic aSyn pathology at earlier time points to progressively make way to a growing population of GCIs, as previously reported^15,22^ (see also **Extended Data Fig. 2).** This is supportive of a neuronal-to-oligodendroglial inclusion transfer as in MSA^24,25^.

Seeding of GCIs by 1B was neither dependent on the mouse strain nor on the intracerebral injection site. When C57BL/6 mice (instead of 129SV) were inoculated with 1B above their right SN, 17 months later, numerous aSyn inclusions could be observed on the right side (**Extended Data Fig. 3a, b**). Incidentally, note the depletion of tyrosine hydroxylase-positive neurons also on the right side in the *pars compacta* of this animal (compare **Extended Data Fig. 3b and c**). Here also, the aSyn inclusions comprised numerous GCIs as demonstrated by co-staining the OL nuclei with Sox10 (**Extended Data Fig. 3a, b, d, e**). They were particularly obvious at the boundary in between the *pars reticulata* and the *pars compacta* and were absent on the left side (**Extended Data Fig. 3a, c, d, f**). Iba1-positive microglial cells were also observed but without evident signs of clustering around aSyn inclusions (**Extended Data Fig. 3d, e, f**).

## Ultrastructure of the neuropathology

Correlative light and electron microscopy (CLEM)^26,27^ was applied to study inclusions in mouse brains (C57BL/6 mice). Consecutive brain tissue slices were imaged by fluorescence microscopy to identify pS129Syn-positive inclusions. Identified sections were then resin-embedded and heavy-metal stained, and serial ultrathin sections were collected consecutively on glass slides and on EM grids. Glass slides were used for immunohistochemistry (IHC) to re-localize the pS129Syn-positive cells and stained with toluidine blue. Recorded images were correlated to EM images taken on the consecutive grids. We used an animal of the 1B-injected series corresponding to **Extended Data Fig. 3a-f**. We imaged 30 pS129Syn-positive inclusions in a region straddling the right GP and CP and followed them throughout the brain tissue by serial sectioning. We could identify fibrillar material and membranous cytoplasmic sub-compartments in the pS129Syn-positive areas. Room-temperature electron tomography on various positions within the IHC-positive regions was used for verification of fibrillar morphology, which showed fibrillar morphology in a dispersed, bundled, and clustered arrangement (**Fig. 1g, Extended Data Fig. 4**). We additionally verified that the fibrillar material in the pS129Syn-positive IHC regions corresponded to aSyn protein assemblies, using immunogold labelling (**Extended Data Fig 5**).

The dispersed fibrillar material correlated with the pS129Syn-positive signal in the IHC image and could be found throughout all observed inclusions in the tissue. Six cells with nuclei of more than 8.5 µm diameter were assumed to be neurons with NCIs, and sixteen pathological cells with a nucleus size below 8.5 µm were assumed to be GCIs in OLs. In many cases, the correlated area consisted of dispersed fibrillar material within a membranous local environment with various vesicular compartments. Fibril bundles could not be observed in all inclusions. If they were present, they varied strongly in length, thickness, curvature, and localization within the cell. Measurement of the identified bundles in 23 inclusions throughout 5 brain slices showed an averaged bundle length of 0.81 ± 0.41 µm. In most cases, the bundle length was of similar size of around 0.62 ± 0.07 µm. In a few exceptional cases, we measured bundles with individual lengths of 1.08 µm up to 3.64 µm (**Fig. 1g**). Fibrils appeared also in bigger clusters in irregular, chaotic arrangements, with fibrils radiating outward (**Extended Data Fig. 4a**).

In addition, we could observe the regular proximity or direct contact of different and morphologically distinct cells to pathological ones (**Extended Data Fig. 6**). The morphology of these cells is reminiscent of cells described as “dark microglia” in mouse models for Alzheimer’s disease^28,29^, as well as of cells recently described in human post-mortem MSA brain^30^.

To further challenge the pathogenic impact of 1B using a “humanized” *in vivo* model, we inoculated 1B fibrils into hemizygous transgenic M83 (M83+/-) mice expressing human A53T aSyn. M83+/-mice have a normal lifespan and do not spontaneously develop fibrillar aSyn inclusions, despite the presence of one copy of the human aSyn A53T transgene^9,10^. We found that the intracerebral injection of 1B fibrils in M83+/-mice systematically caused their rapid demise in a notably synchronized fashion after only 16 weeks (**Fig. 2a**). Animals at death presented numerous intracerebral aSyn inclusions, in particular in the posterior limb of the AC, in which the presence of Lewy neurite bundles, GCIs and GNIs could be observed (**Fig. 2b, c**). Non-seeded M83+/-mice of similar age were healthy and showed no intracerebral aSyn inclusions (**Fig. 2b**). This indicated that like in the two previous wild-type mice strains, 1B fibrils were able to also cause the buildup of an inclusion pathology with cytopathological elements typical of MSA in transgenic mice expressing human A53T aSyn.

**Fig. 2.**
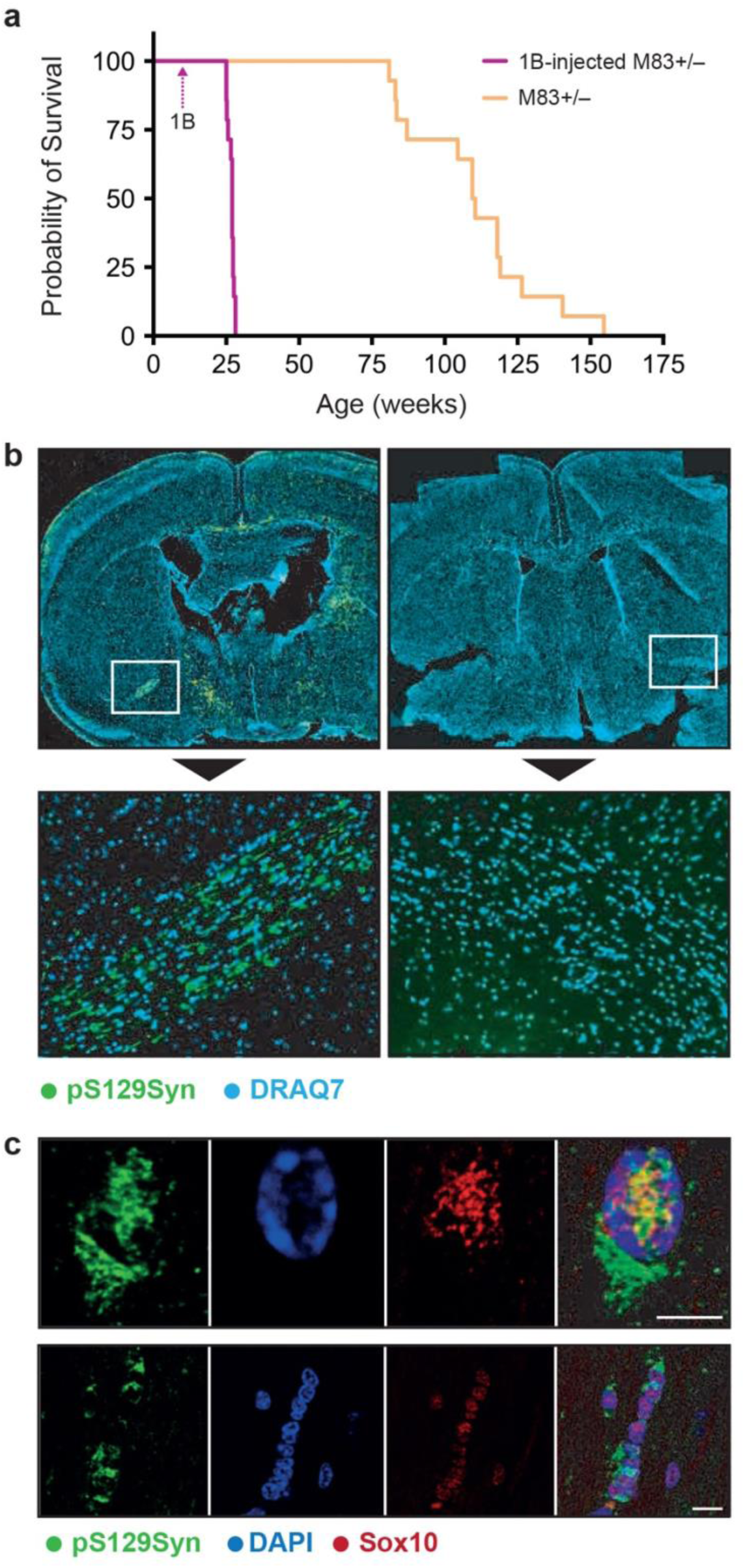
1B is rapidly lethal to humanized M83 +/-mice in which it seeds GCIs and GNIs. **a.** Kaplan-Meier survival curves of 28 hemizygous transgenic mice expressing human A53T aSyn (M83 +/-) without (14 mice, orange curve) of after (14 mice, purple curve) a unilateral injection (right CP) of 1B fibrils (time of injection: arrow). **b:** Illustrative concatenated immunofluorescence views of coronal brain sections of a 1B-injected M83 +/-mouse at the time of death (left), and of a M83 +/-mouse of similar age (right). White boxes: zoomed views centered on the posterior limbs of the AC. aSyn inclusions are revealed using the anti-phospho aSyn antibody EP1536Y (green), nuclei are revealed with DRAQ7 (blue). Note the numerous GCIs and LNs in the AC of the 1B-injected M83 +/-mouse (*nota bene*: the AC is a white matter tract that does not contain neuronal somata). Scale bars = 50 µm. **c:** 1B fibrils seed the buildup of aSyn GCIs and GNIs in oligodendrocytes (OLs) of 83+/-mice. Confocal imaging of a double immunofluorescence of aSyn inclusions (pS129aSyn) and of the OL-specific nuclear marker Sox10 in sections of the anterior commissure. At time of death (4 months post-injection) many commissural OLs harbor GCIs (lower image row). OLs containing both a GCI and a Glial Intranuclear Inclusion (GNII) can also be observed (upper image row). Scale bars = 5 μm.

## 1B and its *in vivo* seeding product 1B^P^

Using cryo-EM, we determined the 1B fibril structure at 1.94Å resolution (**Fig. 3a, Extended Data Fig. 7**). Fibrils were composed of two identical, twisting amyloid protofilaments in 2_1_ screw symmetry, each comprising a pseudo-Greek-key motif consisting of 9 beta-sheets. The map resolved residues G36 to K97, and showed additional density attached alongside the starting sequence from G36 to V40 and reaching toward the inside of the protofilament interface. As that density could not be unambiguously assigned to any sidechains, we did not build an amino acid sequence for this stretch. The protofilaments form their longest beta-sheet secondary structure from amino acids V52 to K58, which forms the site of the main protofilament-protofilament interaction, stabilizing the protofibril interaction by hydrophobic and electrostatic interactions. The main electrostatic clusters are a positive group formed by K43, K45, and H50, and a negative group formed by E57 and T59, flanking the beta-sheet (**Extended Data Fig. 7c**). Additionally, H50 appears to make an inter-rung connection within the same protofilament to the following subunit along the Z-direction to K45 (**Fig 3b**). The interaction between H50 and E57 causes the protofilaments to arrange in a half-step of the typical helical rise, by binding the next aSyn subunit of the opposite strand at the 2.36Å and a resulting twist of 179.52 degrees.

**Fig. 3.**
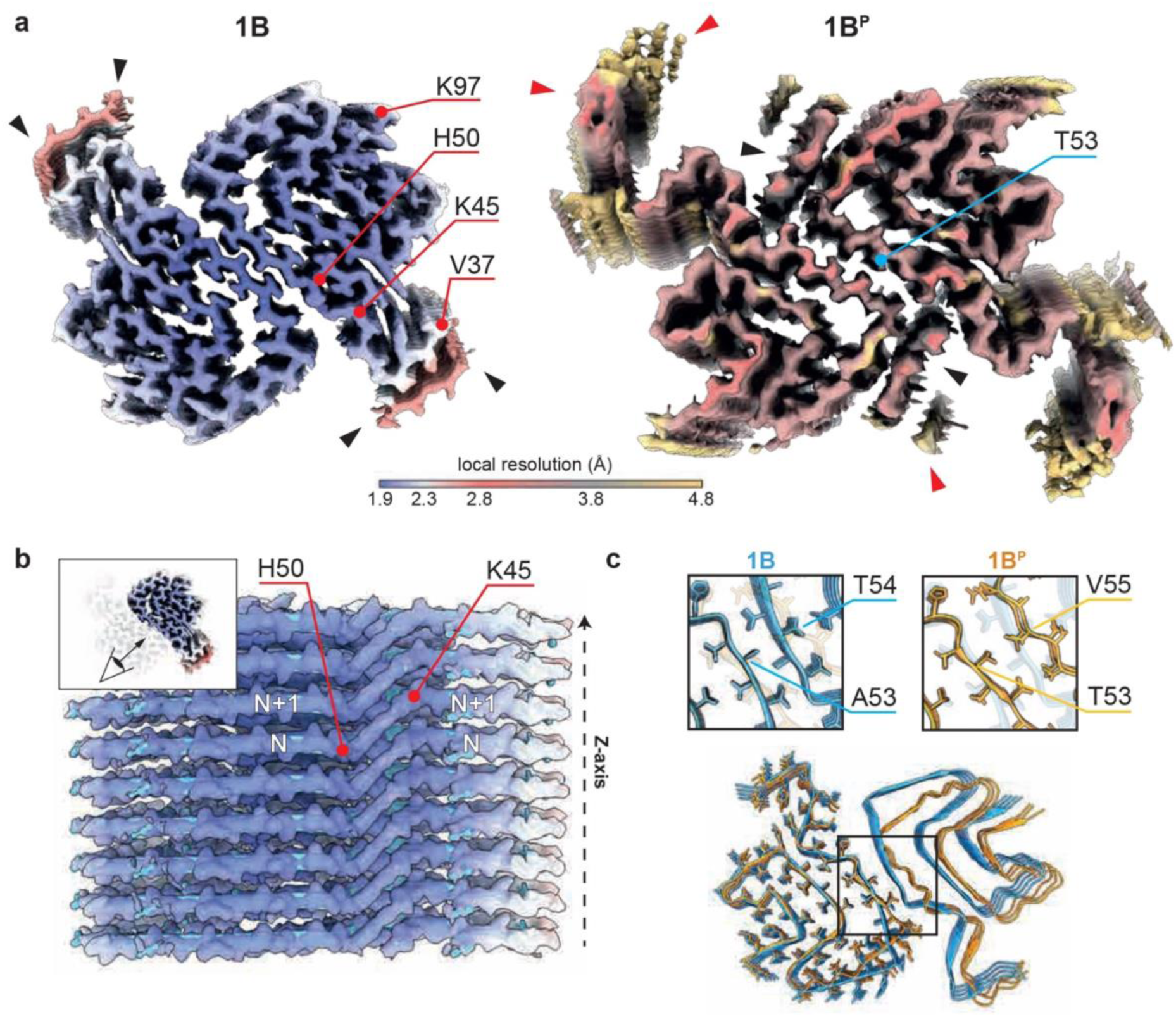
Structural organization of the 1B aSyn fibrils and of the 1B^P^ fibrils seeded by 1B *in vivo*. **a.** Density maps of 1B and 1B^P^ respectively, visualized with the local resolution range from 1.9 to 4.8Å. The model of 1B was built within the resolved densities starting from V37 to K97 (indicated). Wrapping around the stretch starting with V37, an additional density is present on both protofilaments, though at lower resolution (black arrow heads), which can also be observed at different positions in 1B^P^. In addition, the structure of 1B^P^ indicates highly flexible areas that are poorly resolved at those densities and close to the N-terminal region (red arrow heads). **b.** Side view onto the protofilament interface (only one protofilament is shown), revealing the characteristic rise of the amyloid filament and the inter-rung connection at H50 to K47 of the next aSyn subunit (N and N+1). **c.** Sequence overlay of 1B (blue) and 1B^P^ (orange). One protofilament with stick representation, one with only the backbone. The overlay indicates slight alterations in the overall interface of the fibrils, with an RMSD of 1.433Å considering Cα atoms of a single protofilament (see Supplementary Information Table 1). Zoomed views of the protofilament interface highlight changes in interaction between side chains.

To identify the conformation of aSyn fibrils in the seeded pathology we used the fibril purification method previously developed for human MSA brain tissue^11^. The purification was successful in M83 +/-mice, diseased 16 weeks after inoculation with 1B. Despite the low yield of material after the extraction procedure, pS129-aSyn positive fibrils were observed in negative stain imaging (**Extended Data Fig. 8a**). Subsequently, the sample was vitrified for structure determination with cryo-EM.

The extracted material was heterogeneous, with limited concentration of suitable aSyn fibrils for structure determination. We solved the structure of the extracted fibrils, here termed 1B^P^ out of 40 micrographs (∼0.1%) to a resolution of 3.8Å (**Fig. 3a**). 1B^P^ fibrils are composed of two identical protofilaments whose fold is highly similar to the 1B fibril seeding them (overall RMSD of one protofilament: 1.433Å), confirming a prion-like templated misfolding mechanism that underlies inclusion formation in this mouse model. At the same time, an instance of strain adaptation can be observed at the inter-protofilament interface, which spans from V52 to K58 in 1B but is shifted by one amino acid in 1B^P^ where it spans from G51 to E57. This shift, which we suppose is driven by the A53T mutation expressed by M83+/– mice, brings side chains in the 1B^P^ steric zipper that point towards each other into adjacent positions – as in the case of T53 and V55 (**Fig. 3c**) – increasing the distance between Cα atoms in the backbone of the two protofilaments by more than 2Å compared to 1B. As a result, the intercalation of small hydrophobic side chains observed in 1B is lost in 1B^P^, but hydrophobic interactions between opposing protofilaments is still maintained.

The fundamental organization of a single protofilament of 1B not only shows similarities to 1B^P^ but to already published structures as well, mainly the patient-purified MSA fibrils^11^ (PDB: 6XYQ), the corresponding amplification^12^ (PDB: 7NCK), the H50Q variant^31^ (PDB: 6PEO), and wt-and 1-121 aSyn variants^31,32^ (PDB: 80QI and 6H6B) (**Fig. 4, Supplementary Information Table 1**). The superimposition of these structures to 1B highlights differences in the positioning of I88 and F94. The region, previously reported as the C-pocket and acting as the binding site for ThT^33^, is identical in arrangement for a subset of structures and particularly in MSA fibrils (PDB: 6XYQ and PDB: 7NCK) and the H50Q variant (PDB: 6PEO). This is notable because like the 1B fibrils, also the H50Q fibrils (PDB: 6PEO) are ThT negative^31^. Consequently, the inaccessibility of the I88 pocket seems important to account for the ThT-negativity of 1B fibrils^13,14^ (**Fig. 4a**).

**Fig. 4.**
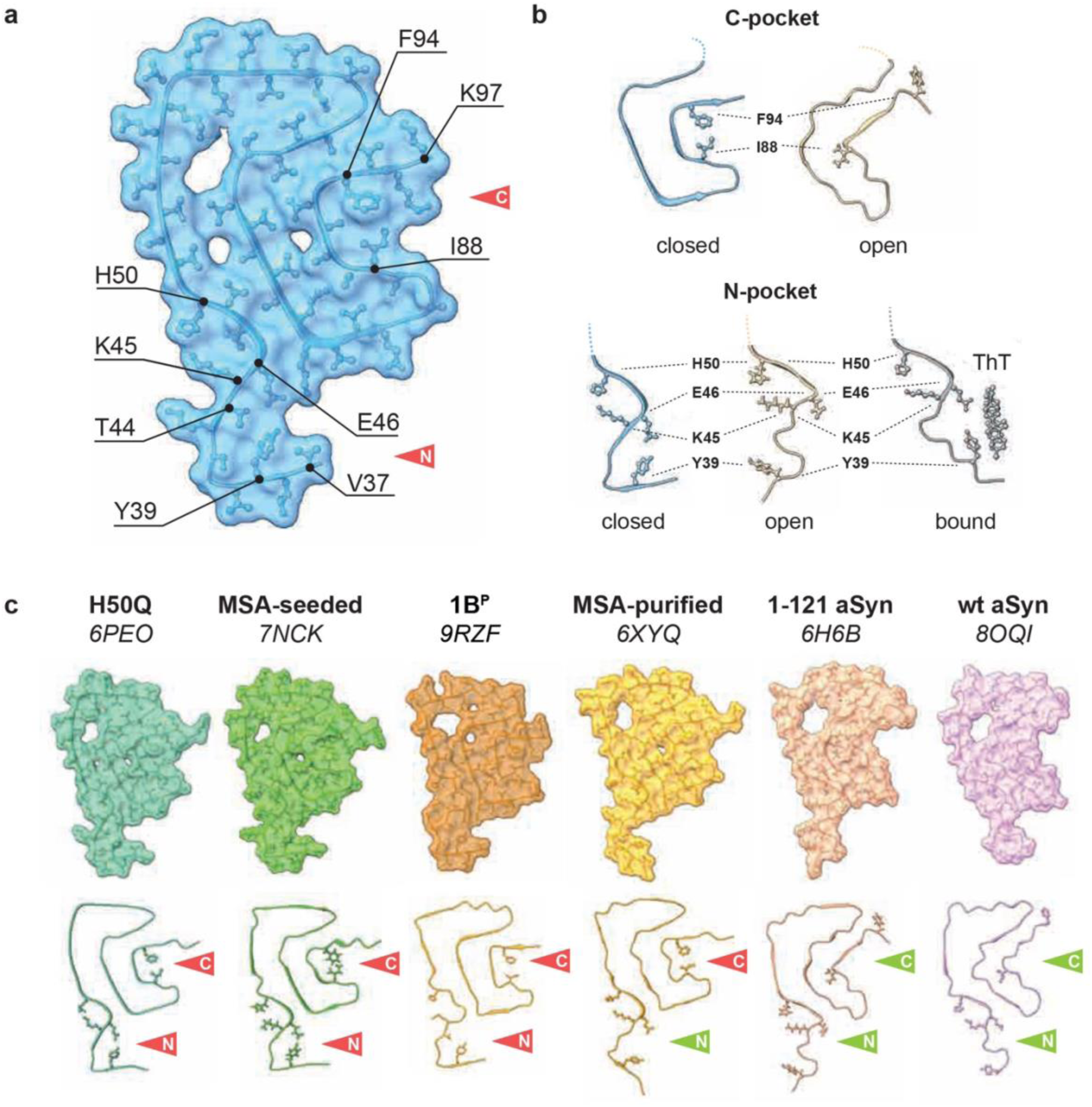
The C and N pockets are the best templated regions between 1B and 1B^P^, are regions where ThT would have got bound (but here they are closed) and are the regions which are also closed in MSA and amplified MSA which is also ThT negative. **a**. Conformation of one protofilament of 1B with surface volume representation. Structural comparisons of single protofilaments with similar fold are summarized with the respective RMSD values in Supplementary Information Table 1. **b.** Zoomed view of the arrangement of the C-pocket on the example of 1B (left, PDB 9EUU) and 1-121 aSyn (right, PDB 6H6B) and the arrangement of the N-pocket on the example of 1B (left, PDB 9EUU), 1-121 aSyn (center, PDB 6H6B), and ThT-bound (grey) aSyn (right, PDB 7YNM). **c.** Shown are the full modeled sequences with transparent surface volume, as well as a ribbon representation with stick representation of Y39, K45, E46, H50, I88, and F94 for all compared structures. The open or closed state of either the C-or N-pocket are indicated by the green and red arrows respectively. Structural similarities are also given as RMSD values in Supplementary Information Table 1.

In addition, the other ThT binding pocket located at the N-terminus (N-pocket)^33^ of the fibrils also shows a modification compared to the structure of other ThT-positive fibrils. In the 1B structure, Y39 is shifted so that it faces T44 and E46. As a consequence, Y39 is not available for interaction with ThT in the N-pocket. Note that a similar conformation of the N-pocket with “shifted” Y39 presumably preventing ThT binding is also observed in the amplified MSA fibrils (PDB: 7NCK)^12^ and the H50Q mutation (PDB: 6PEO), again in line with the parallel observation that seed amplification products templated on MSA brain extracts or on MSA cerebrospinal fluid are ThT-negative^34^.

Collectively, these data support the hypothesis that closure of the ThT-binding pockets at the surface of the fibrils can explain the phenomenon of “ThT invisibility” and at the same time defines the pathophysiological specificity of 1B, 1B^P^ and MSA fibrils.

Besides regions involved in ThT binding, comparison with the other aSyn fibril folds also indicates the near identical arrangement of the H50Q variant to the 1B structure, with the exception of the H50 inter-rung side-chain orientation due to the point mutation. Whether this orientation is important and confers specific “MSA-like” properties to 1B is unknown, especially since the >3Å resolutions of PDB: 6XYQ and PDB: 7NCK do not allow conclusions on its existence in the MSA fibrils.

The 1B fibril structure also shows an outer, non-identified density stretch interacting with G36 to V40 (**Fig. 3a**), a feature which is also present in fibrils amplified from MSA (PDB: 7NCK), in H50Q fibrils (PDB:6PEO), and in 1B^P^ fibrils where poorly resolved densities flank the N terminal (**Fig. 3a**). This is remarkable, because these outer stretches in 1B block access of the disaggregase complex effector-component HSC70 to its binding site at the N-terminus of the fibrils, thus most probably preventing their disassembly and clearance^35^. This structural feature might thus represent a reason for the acute *in vivo* seeding capability of 1B^14^.

## *In situ* pathology characterization

Despite repeated attempts, we were not able to recover enough fibrils from the brains of wild-type mice seeded with 1B to bring them to cryo-EM analysis, most likely because the inclusions are much less numerous than in 1B-injected M83+/-mice. We thus used the conformation-specific amyloid probe h-FTAA^36^ to characterize the GCIs seeded by 1B directly in wild type mouse brain sections without extraction and to also compare them with the inclusions present in MSA (and DLB) post-mortem brain tissue sections. Indeed, and in agreement with previous work^37^ we observed a clear cut difference between the signature of GCIs in MSA and that of Lewy bodies in DLB using fluorescence lifetime imaging microscopy (FLIM) of h-FTAA (**Extended Data Fig. 9**). Using this *in situ* approach we observed that the signature of the MSA inclusions was similar to that of the 1B brain inoculum in our mouse experiments and even closer to the signature of the inclusions seeded by 1B (**Extended Data Fig. 10, Supplementary Information Table 3**). This was also in good agreement with the fact that the inclusion pathology seeded by 1B appeared positive to the MSA-specific silver impregnation staining technique of Gallyas^38^ further suggestive of a structural analogy between 1B, 1B-seeded fibrils in wild type animals, and MSA fibrils in GCIs in human brain tissue sections (**Extended Data Fig. 11**). This is the same conclusion we drew from the cryo-EM structural comparisons of 1B with 1B^P^ and MSA fibrils which were extracted from brain tissue (**Fig. 4, Supplementary Information Table 1**).

## In vivo passage

Finally, given the remarkable similarity between 1B, 1B^P^, and MSA filaments, we sought to determine whether 1B^P^, raised in transgenic animals, could still trigger MSA-like neuropathology in wild-type animals, as synthetic 1B does. To address this question, we faced the challenge of the very low yield of the sarkosyl-based purification protocol for extracting 1B^P^ from 1B-seeded diseased mice—a yield insufficient to obtain the amount of intact fibrils required for *in vivo* reinoculation experiments. To overcome this limitation, we instead used a 10-fold dilution of a total brain homogenate from a 1B-seeded diseased mouse as the inoculum, without any purification step. We also focused on early post-injection effects (6 weeks) to maximize the likelihood of detecting a seeded synucleinopathy driven by trace amounts of 1B^P^ fibrils delivered by the inoculum (i.e., 2 μL of 10-fold diluted whole-brain homogenate). Our results indicate that the amount of 1B^P^ contained in the inoculate reached a critical value sufficient to seed anew an inclusion pathology in wild-type mice with the formation of NCIs, NIIs, GCIs and GNIs (**Fig. 5**) similar to what is observed when using synthetic 1B fibrils as seeds^13–15^.

**Fig. 5.**
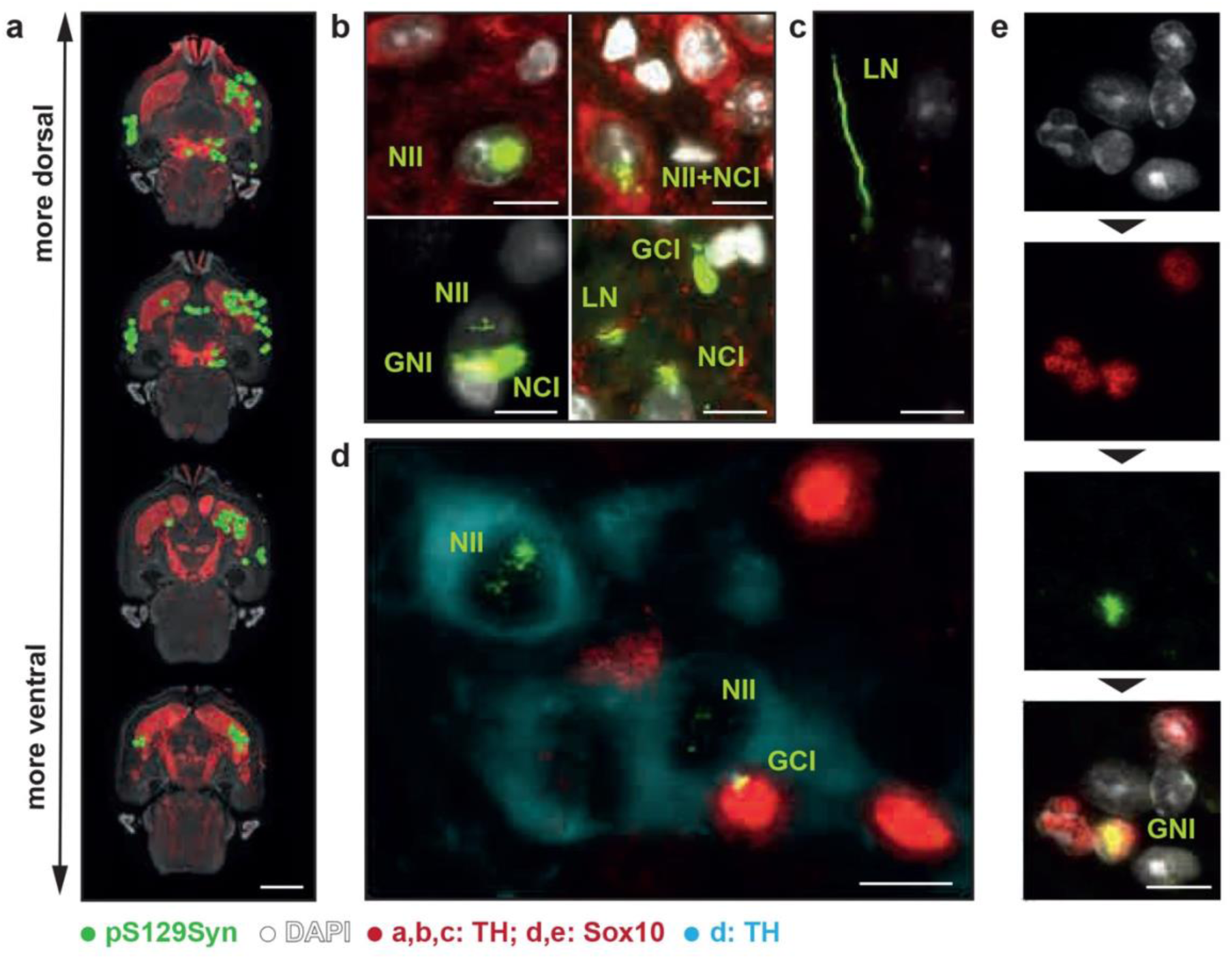
Early seeding of MSA-like pathology in wild-type mice by brain homogenates containing 1BP seeds. Diluted crude brain homogenates were prepared from prematurely dying M83+/-mice injected 4 months earlier with 1B fibrils. These homogenates were injected into the right caudate-putamen (CP) of wild-type 129SV mice. After 6 weeks, mice were sacrificed, and brains were processed for horizontal sectioning and immunofluorescence neuropathology. **a–c**: aSyn inclusions were revealed with anti-phospho aSyn (EP1536Y, green), the nigrostriatal tract with anti-Tyrosine Hydroxylase (TH, red), and nuclei with DAPI (white). **a:** One animal is shown with four sections spanning the nigrostriatal tract dorsoventrally. Green dots annotate aSyn inclusions (Lewy neurites, NCIs, NIIs, GCIs, and GNIs). **d,e:** Examples of MSA-specific aSyn inclusions seeded by 1B^P^-containing homogenates. Sections were stained with TH (turquoise), Sox10 (red, OL marker), pS129Syn (#64, green), and DAPI (white). Images are from the right substantia nigra. **d:** Four dopaminergic neurons (TH-positive cytoplasms) and four OLs (Sox10-positive nuclei) are visible (DAPI omitted for clarity). Three MSA-typical inclusions are present: two neuronal intranuclear inclusions (NIIs) and one glial nuclear inclusion (GNI). **e:** Among eight cells, four OLs aligned in a row and one solitary OL are visible; one OL harbors a nuclear inclusion (pS129Syn, GNI).As control, diluted homogenates from spontaneously dying homozygous M83+/+ mice were used. Although these brains contain aSyn inclusions, they are not MSA-type, and their homogenates failed to induce inclusion pathology in recipient wild-type mice (not shown).Scale bars: 3 mm **(a)**, 10 µm **(b–e)**. 3D reconstructions and corresponding movies of selected inclusions are provided in **Supplementary Information Videos 2–6**.

Given the structural similarity of the synthetic 1B seed - which does not undergo any sarkosyl purification before cryo-EM analysis - and of 1B^P^ which does go through this detergent-based purification step before cryo-EM, we can deduce that the extraction procedure has no significant impact on the structural arrangement of 1B^P^ which is essentially acquired by templating. In addition, and in line with the notion that “non extracted” 1B^P^ and 1B have similar structures, the pathology seeded by brain homogenates containing 1B^P^ is identical to the one seeded by synthetic 1B.

Altogether, this indicates that the 1B strain characteristics are encoded in the pseudo-Greek-key motif of the 1B/1B^P^ fold rather than in the unattributable densities specifically observed in either 1B or in 1B^P^ after adaptation. It also indicates that the widening and the sliding of the protofilament interface acquired in 1B^P^ do not impinge on the MSA-like pathology-inducing characteristics of 1B/1B^P^.

This is in agreement with the observation that the seeding of inclusions by 1B in cultured neurons is insensitive to mutations affecting residues located at the protofilament interface (H50Q, G51D, A53T and A53E) but that instead, it is abrogated by the mutation E46K which disrupts the E46-K80 salt bridge ‘latch” necessary for the stabilization of the pseudo-Greek-key motif in the 1B fold (Extended data Fig. 12). Notably, the differential impact of the various aSyn mutations on the replication of 1B we report here is similar to what was reported for so-called MSA “prions” by Prusiner and his group^39,40^.

In conclusion, the unattributed densities in 1B and 1B^P^, as well as the interface adaptations observed in 1B^P^, do not appear to play a significant role in the pathogenic activity of 1B/1B^P^. In other words, the fold kernel shared by 1B, 1B^P^, and MSA filaments is likely necessary and sufficient to induce both MSA and MSA-like neuropathologies.

The fold kernel of 1B, which is templated with the highest fidelity in 1B^P^, forms the closed N- and C-pockets (**Supplementary Information Table 1**). As previously discussed, such closure accounts for the ThT-invisibility of both 1B^13,14^ and the amplified MSA filaments^34^. This is particularly relevant given that ThT-invisibility is also observed in two other synthetic fibril assemblies of unknown structure, both of which were reported to seed GCIs *in vivo*^21,22,41^.

It is thus tempting to speculate that closed N- and C-pockets are also present in these latter fibrils, further supporting the idea that, within the fold kernel shared by 1B, 1B^P^, and MSA filaments, the closure of the N- and C-pockets represents the minimal common structural determinant responsible for both MSA and MSA-like pathologies induction.

## Discussion

We present the structures of aSyn fibrils 1B as well as of their *in vivo* seeding product 1B^P^. Indicative of a high-fidelity conformational replication process, the structures present a remarkable similarity and are capable of perpetuating MSA-like pathology through passage in humanized and wild-type mice.

This study provides evidence of a protein-only replication process occurring *in vivo* and evidenced at a 3.8Å resolution. The templating fidelity of the synthetic 1B seeds, both *in vitro* and in the mouse brain, is remarkable given the difficulty to efficiently amplify MSA brain extracts with structural preservation^12^. While we cannot rule out the presence of additional fibril polymorphs different from 1B^P^ in the 1B-inoculated mouse model, which may have arisen through secondary nucleation and are undetectable by cryo-EM, the robust propagation of the 1B protofilament fold provides strong support for a prion-like templating mechanism.

This notion is strengthened by our observation that brain homogenates containing 1B^P^ can trigger in wild-type mice the same MSA inclusion pathology as the one seen after 1B injection, indicating that untemplated secondary nucleations, if any, do not significantly participate in the MSA pathology seeded by 1B. Instead, the kernel common to 1B and 1B^P^ and here termed “MSA kernel” defines through a templated replication the strain characteristics of the fibrils. This MSA kernel is shared in one of the two protofilaments of MSA fibrils extracted from patient 5 in the seminal study of Schweighauser and colleagues^11^, as well as the MSA fibrils resulting from the *in vitro* amplification of the same patient’s sample^12^. Thus far, it was not possible to purify the fibrils naturally occurring in homozygous M83 +/+ to compare them with the templated structure of 1B^P^ using cryo-EM. However, we observed that brain homogenates from dying M83 +/+ mice were not capable of eliciting any synucleinopathy when injected into the brain of wild-type mice like it is the case for 1B^P^-containing brain homogenates (**Fig. 5**). This suggests that either the fibrils in the brains of spontaneously dying M83 +/+ mice are a different polymorph, or the fibril titer is too low for stable transmission. Interestingly, the templating fidelity of the MSA kernel between 1B and 1B^P^ is most pronounced in two particular regions: the N- and C-pockets. These 2 regions are of particular interest because they define the binding sites for the amyloid probe ThT^33^, but are closed and not accessible to the dye in the MSA kernel. Our data suggest that structural features of aSyn fibrils – specifically closed N- and C-pockets – are related to their ThT invisibility and their seeding properties, in particular their ability to seed GCIs. This is in agreement with the fact that the 3 synthetic fibril assemblies that have been reported to induce GCIs in mice and rats are all weak ThT binders, and that the fibrils amplified from MSA patient brain samples and CSFs^34^ are ThT negative as well.

This collectively points to a major pathophysiological role of C- and N-pocket closure with direct relevance to MSA.

In the MSA kernel, the specific binding site around the C-pocket of I88 is closed by forming a “lever” with F94, mainly caused by the outward orientation of I88. Compared to other aSyn fibril structures with an open C pocket, in 1B/1B^P^, the inner “hook” of the pseudo-Greek-key motif along the stretches of V74 to F94 is shifted by one amino acid position starting at A85. As a result, A85 points outwards at the most prominent position of the inner hook, which causes a consequential inversion of sidechain positioning for the remaining sequence. In 1B/1B^P^, this causes S87, a known modification site for fibril inhibition by O-Glc-N acylation^42^ or by phosphorylation^43^ to be inaccessible, and I88 to be positioned outwards, undergoing a connection with F94 and therefore closing the C-pocket.

Enforcing the post-translational modification of S87 or “forcing open” the C-pocket using small molecules might thus represent promising means to derail the templating process during fibril growth, and force fibril elongation to take place on a modified template which has lost part of the MSA kernel. This would presumably not prevent fibril growth but could perhaps impinge on the acute replicative and toxic properties of MSA fibrils and slow down disease progression.

The closed N-pocket is a shared characteristic of the 1B and 1B^P^ structures, and this was also observed in the amplified MSA filaments from patient 5 ^12^. However, fibrils extracted from MSA patients^11^ did not share the closed N-pocket conformation (**Fig. 4, Supplementary Information Table 1**). Whether this points to a dispensable pathogenic role of N-pocket closure remains to be determined.

Finally, we note that the ability of crude brain homogenates from MSA patients to seed pathological inclusions in wild-type mice has – to our knowledge – not been reported to date. This may be due to a lack of systematic investigation, or it may suggest that the fibrils present in human MSA brain samples are intrinsically less aggressive than 1B or 1B^P^ fibrils. This latter would explain why human MSA brain extracts trigger little MSA-like pathology in M83+/-animals^9^. If so, the higher infectivity of the 1B/1B^P^ strain could result from the specific structural features not shared with patient-derived fibrils. Notably, these include: (i) a closed versus open N-pocket; (ii) the pseudo-symmetric arrangement of two identical protofilaments in 1B/1B^P^, as opposed to the asymmetric fold of two distinct protofilaments in MSA fibrils; and (iii) additional, unattributed densities in 1B and 1B^P^, which — although distinct — may obscure chaperone clearance sites in both. The relevance of these structural differences remains to be determined. Regardless of the precise mechanism underlying the high seeding efficiency, it is important to emphasize that, in the absence of prior structural knowledge of synthetic or patient-derived α-synuclein fibrils, such materials should be handled with extreme caution.

Taken together, this study demonstrates the existence of a synthetic aSyn fibril strain that is capable of inducing MSA-like pathology in wild-type mice and replicates in M83+/-mice through a protein-only templating process. The structures of the fibril seeds and of fibrils formed *in vivo* highlight the similarity of 1B and MSA fibrils, and postulate a link between fibrillar structure, assembly, mechanism of prion-like seeding and spreading, as well as animal-to-animal transmission.

## Methods

### aSyn expression

*Escherichia coli* strain BL21(DE3) was transformed with pT7-7 aSyn *wt* vector^43^ from Addgene, (plasmid #36046; http://n2t.net/addgene:36046; RRID: Addgene_36046) and plated onto a Luria broth (LB) agar plate containing ampicillin (100 μg/mL) and 1 g/L glucose. A preculture in 5 mL of LB medium was inoculated with one clone and incubated at 37°C under 200 rpm shaking for 4 h. Cells from the LB preculture were recovered by centrifugation (1000× g, 10 min) and used for inoculating 200 mL of LB medium. Cells were grown overnight at 37°C under 200 rpm shaking and then diluted in 2 L of culture. Protein expression was induced by adding 0.5 mM isopropyl-β-d-thiogalactopyranoside during the exponential phase, evaluated at an optical density at 600 nm reaching 0.6. Cells were harvested after 5 h of culture at 30°C by a 7000× g centrifugation at 4°C, 20 minutes (JLA 8.1000, Beckman Coulter, Villepinte, France), and pellet was kept at −20 °C until purification.

### aSyn purification

The pellets were resuspended in lysis buffer (10 mM Tris and 1 mM EDTA (pH 7.2) with protease inhibitor tablets (cOmplete, EDTA-free protease inhibitor cocktail, Roche, Basel, Switzerland) and sonicated at 50% max energy, 30 s on and 30 s off for three rounds with a probe sonicator (Q-Sonica, Newtown, CT, USA). The sonicated pellets were centrifuged at 20,000× g for 30 min, and the supernatant was saved. The pH of the supernatant was then reduced to pH 3.5 using HCl, and the mixture stirred at room temperature (RT) for 20 min and then centrifuged at 60,000× g for 30 min. The pellets were discarded. The pH of the supernatant was then increased to pH 7.4 with NaOH and then dialyzed against 20 mM tris-HCl (pH 7.4) and 100 mM NaCl buffer before loading onto a 75 pg HiLoad 26/600 Superdex column equilibrated with the same buffer with ÄKTA pure system. Monomeric fractions were collected and concentrated if needed by using Vivaspin 15R 2 kDa cutoff concentrator (Sartorius Stedim, Göttingen, Germany). Purification fractions were analyzed by polyacrylamide gel electrophoresis (PAGE) tris-tricine 13% dying with ProBlue Safe Stain. Protein concentration was evaluated spectrophotometrically by using absorbance at 280 nm and an extinction coefficient of 5960 M^−1^ cm^−1^. Quantification of the preparations with a Pierce chromogenic LAL kit indicated a low endotoxin [lipopolysaccharide (LPS)] residual value of 0.03 to 0.06 EU per μg of recombinant protein.

### aSyn fibrillization

Solutions of monomeric aSyn at 4 to 5 mg/mL in saline (H2O, 100 mM NaCl, and 20 mM tris-HCl pH 7.40) were sterilized by filtration through 0.22 μm Millipore single-use filters and stored in sterile 15 mL conical falcon tubes at 4°C. Sterilized stock was then distributed into safe-lock Biopur individually sterile-packaged 1.5 mL Eppendorf tubes as 500 μL aliquots and were seeded with 1% of 1B strain fibrils^13–15^. The tubes were cap-locked and additionally sealed with parafilm. All previous steps were performed aseptically in a particle-free environment under a microbiological safety laminar flow hood. The samples were loaded in a ThermoMixer (Eppendorf, Hamburg, Germany) in a 24-position 1.5 mL Eppendorf tube holder equipped with a heating lid. The temperature was set to 37 °C, and continuous shaking at 2000 rpm proceeded for 4 days. 1B templating of the fibrillization products was quality-checked using the fibrilloscope^13–15^.

### Sonication

Prior to intracerebral injections, 1B aSyn fibril stocks (4 mg/mL) were distributed in cap-locked, sterile 0.5 mL polymerase chain reaction (PCR) tubes (Thermo Fisher Scientific, Bordeaux, France). Sonication was performed at 25 °C in a Bioruptor Plus water bath sonicator (Diagenode, Liège, Belgium) equipped with thermostatic control and automated tube carousel rotator. The sonication power was set to “high”, and 10 cycles of 30 s on followed by 10 s off were applied. In agreement with our previous quantifications^14^, over 80% of the fibrils were 50 nm-long or less after application of this protocol (not shown).

### *In vivo* aSyn pathology

Adult 129SV (intrastriatal injections), C57BL/6 (nigral injections), and transgenic M83 (hemizygous, intrastriatal injections) were housed in a temperature-controlled (22 °C) and light-controlled environment on a 12 h light/12 h dark cycle with forced ventilation (humidity below 50%) and with access to food and water ad libitum. All experimental procedures were conducted in accordance with the European Communities Council Directive (2010/63/EU) for care of laboratory animals. The study design was approved by the ethics committees of the University of Bordeaux, of the ANSES/Ecole Nationale Vétérinaire d’Alfort/Université Paris-Est Créteil, and of the University of Salento, and authorized by the French Ministry of Higher Education and Research and by the Italian Ministry of Health (APAFIS #33147-2021091711598830 v6, APAFIS #37712-2022061615206629 v8, 0013178-P-17/05/2019). The mice (6–8 weeks old) unilaterally received 2 μl of sonicated aSyn fibrils 1B (4 mg/mL) by stereotactic delivery at a flow rate of 0.4 µl/min, and the pipette was left in place for 5 min after injection to avoid leakage. Delivery was performed within the right striatum (coordinates from bregma: AP, −0.1; L, +2.5; DV, +3.8), or above the right *substantia nigra* (coordinates from bregma: AP, −2.9; L, −1.3; DV, −4.5). Wild-type mice (n=20) and transgenic M83 hemizygous mice (n = 28) injected with either PBS, 1B fibrils, or brain homogenates were euthanized after 6 weeks, 17 months, 24 months, or when reaching the humane endpoints defined as death regarding the transgenic M83 mice, and were transcardially perfused with either tris-buffered saline (pH = 7.4) followed by 4% paraformaldehyde in PBS pH = 7.4 at 4 °C, or with cacodylate buffer 0.1 M with 1 mM CaCl_2_ (pH = 7.4) followed by 4% paraformaldehyde plus 0.1% glutaraldehyde in 0.1 M cacodylate buffer (pH = 7.4). Brains were subsequently postfixed in the same fixative. For standard neuropathology, the brains were paraffin embedded, and 10 µm sections were cut with a rotative microtome (Leica, Milan, Italy). The sections of interest were deparaffinized and processed for epitope retrieval: the slides were immersed in citrate buffer pH 6 (Dako Agilent Technologies, Les Ulis, France) and placed in a pressure cooker (Bio SB, Santa Barbara, CA, USA) at 114°C-121°C for 10 min. After a cooling period of 20 min, the slides were washed twice for 5 min in PBS at room temperature. They were then processed for simple or double immunofluorescence using the following primary antibodies diluted at 1/500, or their combinations: EP1536Y (Abcam) or pSyn#64 (Wako) for detecting phospho-S129– positive aSyn inclusions; LB509 for detecting human aSyn (Abcam, mouse monoclonal); anti-Sox10 (Abcam, rabbit monoclonal) for detecting OL nuclei; EP1532Y and anti-TH raised in chicken (anti-Tyrosine Hydroxylase, Abcam) for detecting the nigrostriatal tract; and anti-Iba1 (Abcam) for detecting microglia. DRAQ7 or DAPI were used to image the nuclei. The AlexaFluor-coupled secondary antibodies were from Thermo Fisher (Alexa 488, 568, and 674). The sections were acquired using a Pannoramic slide scanner (3D HISTECH, MM France) in epifluorescence mode, and multichannel fluorescence optical sections of the samples were performed (thickness < 0.8 µm) using either a Zeiss CD7 platform of a Leica SP5 Laser Scanning Confocal Microscope equipped with a spectral detector, 488, 561, and 633 nm laser lines, a motorized X-Y stage, and a mixed stepping motor/piezo Z controller. The objective was 40×, and Z step size was set to 0.5 µm to produce stacks of 15 to 20 Z planes. The pinhole was set to 1 airy unit. For 3D reconstructions and volume rendering/animations (corresponding to 360° tilt series of composite max pixel projections images), raw 3-channel Z-stack images were processed offline using Icy^44^(v2.4.0.0).

### High content analysis (HCA)

Timed inbred-pregnant C57BL/6JOlaHsd (aSyn KO) female mice were received from Envigo (Puteaux, France) 2 days before initiation of the primary culture. The pregnant mouse was euthanized by cervical dislocation, and the aSyn KO embryos (embryonic day 18) were surgically extracted and cold-euthanized. Cortices were harvested from 8 of them, pooled, and dissociated enzymatically and mechanically (using a neural tissue dissociation kit, C Tubes, and an Octo Dissociator with heaters; Miltenyi Biotech, Germany) to yield a homogenous cell suspension pool of 8 individuals. The cells were then plated at 20,000 per well in 96-well plates (Corning, BioCoat poly-d-lysine imaging plates) in neuronal medium (MACS Neuro Medium, Miltenyi Biotech, Germany) containing 0.5% penicillin-streptomycin, 0.5 mM alanyl-glutamine, and 2% NeuroBrew supplement (Miltenyi Biotech, Germany). The cultures were maintained with 5% CO2 at 37°C in a humidified atmosphere. The medium was changed by one-third every 3 days, until 30 DIV (days in vitro). In such cultures, and under control conditions, neurons represented approximately 85 to 95% of the cell population; thus, for simplicity, they are here referred to as “neurons.” After 7 DIV, vehicle and sonicated 1B aSyn fibrils were added at a final concentration of 10 nM equivalent monomeric concentration. When relevant, neurons were infected at DIV 10 with AAV (adeno-associated virus) particles (multiplicity of infection, 1000) carrying the cDNA of aSyn and its variants under the control of the human synapsin promoter, as previously described^14^. High Content Analysis was performed after fixation and double immunofluorescence staining with the conformation-dependent SynF1 antibody (mouse monoclonal, BioLegend) and the human aSyn-specific MJFR1 antibody (rabbit monoclonal, Abcam) as previously described on images acquired at 20× using the generic analysis module of an Incucyte S3 (Sartorius, USA) with Top-Hat segmentation^14^. For quantifications the signal integral of the segmented areas in each field of view was recorded. Primary neuronal cultures from C57BL/6 embryos physiologically expressing endogenous mouse aSyn and not transduced with any AAV vector (**Extended data Fig. 1**) were produced and analyzed similarly but after fixation at DIV24 and after revealing pS129Syn with EP1536Y and human aSyn with LB509.

### Correlative light and electron microscopy

The CLEM protocol was performed as previously described^26,27^. In short, 20 to 40 µm thick sections were cut from perfused mouse brain on a vibratome (LEICA VT1200). Immunolabeling against aSyn pathology allowed for the identification of immuno-positive sections to be used for further processing with EM. For this, the sections were incubated overnight with a primary EP1536Y (Abcam) antibody at 4°C followed by Alexa-conjugated secondary antibody staining together with DAPI for cell nuclei identification. Fluorescence images were acquired with a Leica Thunder Tissue Imager equipped with a K8 fluorescent camera. Chosen sections with strong aSyn pathology were prepared for EM by post-fixing in reduced 2% osmium, thiocarbohydrazide and 2% Osmium tetroxide, followed by contrasting in 2% uranyl acetate and lead aspartate treatment at 60°C. The sections were dehydrated with an increasing acetone gradient, before infiltration with EMBED 812 resin for final flat embedding. The hardened sections were imaged again by light microscopy and the sections overlayed with the overview fluorescence map by using identifiable tissue features. Regions of interest were marked and the coordinates used for laser dissection. The resulting blocks were laser cut using a Leica LMD7 and glued onto a resin block. Serial sectioning with an ultra-microtome (Leica UC7) produced 120 to 200 nm thick tissue sections that were alternatingly collected on EM grids (slot and hexagonal grids) or glass slides. Glass slides were further processed for immunohistochemistry and toluidine blue staining.

Light microscopy images were acquired at 40x magnification and correlated to each other for the generation of an image overlay of pathology signal (IHC) and high contrast tissue morphology (toluidine blue). The resulting overlays were then correlated to low magnification (640x nominal magnification) EM overviews for cell identification by using identifiable tissue features both in the LM and EM images. The identified pathology was finally imaged and merged at high magnification in a tile acquisition series. For EM image acquisition, a Thermo Fisher Scientific (TFS) Talos F200C with a CETA camera or a CM100 (Phillips) with a TVIPS F416 camera were used. LM and EM images were adjusted for brightness and contrast where necessary.

### On-grid immunogold labelling and analysis

EM grids with tissue sections were placed on a droplet of etching solution (1% periodic acid in water, Sigma Aldrich, United States), with the sections facing down, for 2.5 minutes. Grids were washed 3 times for 2 minutes in ddH_2_O, then washed 2 times for 2 minutes in washing buffer (stock solution of 10% BSA, Aurion, which was 1/50 diluted in ddH_2_O), and blocked in blocking solution (Aurion) for 5 minutes to reduce non-specific binding. Grids were then incubated with primary antibody (EP1526Y, Abcam, 1/50 diluted in washing buffer) for 60 minutes, washed 6 times for 2 minutes in blocking buffer, and then incubated in a solution of gold nanoparticles with a size of 10 nm (protein A gold, Aurion, 1/50 diluted in washing buffer) for 90 minutes. They were washed 3 times for 2 minutes in blocking solution, 3 times for 2 minutes in washing buffer, and 4 times for 1 minute in ddH_2_O. Finally, grids were post-stained in 1% uranyl acetate for 1 minute and washed 3 times for 10 seconds in ddH_2_O. Sections were imaged with a transmission electron microscope (TFS Talos L120C), operated at an operating voltage of 120 kV, and images were recorded with a TFS Ceta camera.

Gold particles were detected automatically in the recorded EM images, using FIJI^45^, by applying a difference-of-gaussian filter (sigma1 = 10 nm, sigma2 = √2 • sigma1). The resulting image was binarized with a manually selected threshold. Particle detections with an area smaller than 25 nm^2^ and a circularity (4π•area/perimeter^2^) below 0.8 were excluded from analysis. The local density of gold was calculated as the decadic logarithm of the sum of a pixel’s distance to the 15 closest neighboring gold particles. The gold particle detection was implemented as a FIJI macro and the density calculation as a python notebook.

### Electron tomography and 3D reconstruction

Electron tomography data was collected using the Tomography software (TFS) with multi-site batch acquisition on a TFS Talos F200C with tilt series covering -60° to +60° in 2° steps. Tomograms were reconstructed using IMOD^46^ and filtered with a non-local-means filter with Amira (TFS Amira version 2021.2). For segmentation, nuclear membranes, fibrils, and other membranous parts were manually annotated and traced throughout the tilt-series using Amira. Every few slices the respective feature of interest was selected by drawing directly onto the image and then interpolated to generate an accurate 3D volume. The final volume was visualized directly in AMIRA.

### aSyn purification from brain homogenate

Extraction of the sarkosyl-insoluble fraction of a transgenic M83+/-mouse brain 16 weeks after inoculation with 1B was done using the same procedure as was previously published for fibrils purified from a MSA patient^11^. In brief, brain tissue was homogenized in 20 vol. (v/w) extraction buffer consisting of 10 mM Tris-HCl, pH 7.5, 0.8 M NaCl, 10% sucrose and 1 mM EGTA. Homogenates were brought to 2% sarkosyl and incubated for 30 min. at 37° C. Following a 10 min. centrifugation at 10,000 g, the supernatants were spun at 100,000 g for 20 min. The pellets were resuspended in 500 μl/g extraction buffer and centrifuged at 3,000 g for 5 min. The supernatants were diluted 3-fold in 50 mM Tris-HCl, pH 7.5, containing 0.15 M NaCl, 10% sucrose and 0.2% sarkosyl, and spun at 166,000 g for 30 min. Sarkosyl-insoluble pellets were resuspended in 100 μl/g of 30 mM Tris-HCl, pH 7.4.

### Immunogold labeling and negative staining of sarkosyl purification

3 µl of non-diluted sample was applied to 100 mesh Ni grids with formvar support (Electron Microscopy Sciences) that were glow-discharged at 15 mA for 60 seconds. After 10 minutes incubation time, the grid was blotted, washed 3x 5 mins with incubation solution (0.2% BSA-c/PBS; Aurion), and incubated for 1h at room temperature in a solution of aSyn-pS129 antibody (EP1536Y, Abcam, ab51253) diluted 1:100 in incubation solution. The grid was washed 6x 5 min and incubated for 2h in a solution containing gold nanobeads (diameter 6 nm) coupled to an anti-rabbit antibody (Aurion), diluted 1:50 in incubation solution. After washing 6x 5 min in incubation solution and 3x 5min in PBS, the grid was immersed in 1% uranyl acetate for 40 seconds, blotted, and air-dried. The grid was imaged with a Talos transmission electron microscope at an operating voltage of 200 kV (ThermoFisher Scientific).

### Cryo-electron microscopy

For recombinant 1B fibrils: Cryo-EM grids were prepared using a Leica GP2 plunge freezer in a BSL-2 environment at room temperature. Quantifoil 300 mesh gold grids were glow discharged in air for 90s. Grids were prepared using 3 µl of 1:1 and 1:2 diluted fibril samples in preparation buffer (H2O, 100 mM NaCl, and 20 mM tris-HCl. pH 7.40) and vitrification by rapid freezing in liquid ethane. The grids were screened for the presence of non-clustered fibers and high-resolution images were acquired with an automated EPU (v3.10) setup on a TFS Titan Krios G4 (TFS) electron microscope equipped with a cold-FEG 300kV electron source and a Falcon4i camera. 11’879 dose-fractionated movies (EER) were collected with a total dose of 40 e^-^/Å^2^ at a pixel spacing of 0.66 Å at the sample level. A defocus range between -0.8 μm and -1.8 μm was used during the acquisition.

### For purified 1B^P^ fibrils

The sarkosyl-insoluble fraction of a transgenic M83+/-mouse brain was diluted 1:3 in Tris-HCl, 50mM, 150mM NaCl, pH7.4, and 2.5 µl was applied to a Quantifoil 300 mesh gold grid, coated with an additional 5-nm thick carbon layer. The grid was not glow-discharged. 30 seconds after applying the sample, the grid was blotted from the front side for 6 seconds and plunge-frozen at room temperature and 80% humidity in a Leica GP2 plunge freezer. A total of 42,812 exposures was collected on a Titan Krios (TFS) electron microscope equipped with a cold-FEG 300kV electron source and a Falcon4i camera. The total dose applied per movie (EER) was 50 e^-^/Å^2^ and the pixel spacing was 0.732 Å. A defocus range between -0.8 μm and -2.2 μm was used during the acquisition.

### Helical reconstruction and model building of recombinant 1B fibrils

All the subsequent image processing and helical reconstruction was carried out in RELION 4.0^47^ (**Extended Data Fig. 13**). The recorded movies were gain corrected, motion-corrected and dose-weighted using RELION’s own implementation of MOTIONCOR. The raw EER files were dose-fractionated to a total of 32 frames (1.23 e^-^/Å^2^ per frame). After CTF estimation using Ctffind 4.1^48^, the aligned averages that had an estimated CTF resolution of better than 5Å were selected, resulting in a total of 11,879 micrographs. aSyn fibers were picked manually as start-to-end helical segments and extracted with an inter-box distance of 19 Å. Several rounds of reference-free 2D classification were performed to exclude suboptimal 2D segments from further processing. The four best classes were selected and binned 4 times to re-extract the segments with a bigger box size encompassing the entire helical crossover. Clearly visible twisted class averages containing the entire crossover were used to generate initial de novo models using the relion_helix_inimodel2d program. The most likely twist was estimated from a crossover distance measured from 2D class averages and corroborated with the direct measurements from micrographs.

The helical segments (105,496) from the entire dataset were then re-extracted with 600 pixel box size (binned by 2) comprising about 40 % of the crossover. Exclusion of the suboptimal classes after 2D classification resulted in 64’889 particles. Multiple rounds of 3D auto-refinement were carried out to optimize the helical twist and rise, and to check for the correct symmetry operators. This was validated from the reconstructions showing clear separation of beta-strands along the helical axis. Imposing a 2_1_ cork-screw symmetry (pseudo two-fold symmetry) yielded the clear separation of beta-strands when compared to C1 and C2 symmetries. A 3D classification without image alignment was further performed to exclude the segments that gave suboptimal 3D volumes. This resulted in a final 51,272 particles. These were finally re-extracted at a 512 pixel (unbinned) box size and subjected to 3D auto-refinement, followed by CTF refinement. Further iterations of Bayesian Polishing coupled with CTF refinement yielded a map at 1.94 Å. The maps were sharpened using the standard postprocessing procedure in RELION with an ad-hoc B-factor of 21 Å^2^.

The atomic model for the backbone of the core of the 1B fibrils encompassing residues 34 to 95 was built de novo using Coot^49^ (v0.9.8.96) Initially, three beta-rungs were modeled in coot and were refined in real-space in PHENIX^45^ (v1.21.2) Finally, these chains were extended to 9 beta-rungs per protofilament and refined in tandem with phenix.real_space_refine and in coot to improve the Ramachandran and Geometric statistics. NCS and secondary structural restraints were included during the iterative refinement process.

### Helical reconstruction and model building of purified 1BP fibrils

Because most of the 42,812 collected movies did not contain any fibrillar material, they were first imported into cryoSPARC^50^ (v4.7.0), motion corrected, and manually inspected using a “manually_curate_exposures” job. After manual inspection, 156 movies with intact, twisting fibrils were selected for further processing in RELION4^47^. They were dose-fractionated in RELION’s MOTIONCOR implementation with 24 frames per fraction, corresponding to a dose of 1.28 e^-^/Å^2^ per frame, and CTF-corrected using Ctffind 4.1^48^. Fibrils were found to cluster into three morphological groups (**Extended Data Fig. 8**). Manual picking was done in three rounds, using 209, 7 and 40 movies for the three respective groups. After extraction with binning of 2 and an inter-box distance of 14.4 Å (3 asymmetrical units), several rounds of 2D classification were applied. One of the three morphological groups (40 movies) yielded 2D classes for which a clear twist and monomer separation was visible, while the other two morphological groups did not yield any interpretable result.

Using the 2D class averages of the one usable group, a de-novo initial model was generated with relion_helix_inimodel2d. Particles were re-extracted with a box size of 800 and a binning of 2, and after multiple rounds of 2D classification, 1913 particles were kept for a 3D classification job with 1 class and the de-novo inimodel2d reconstruction as initial model. When a fibril backbone was visible, the regularisation parameter (T) was gradually increased to 12 over the course of 50 iterations, and the result was used for a 3D refinement with a tau2 fudge factor of 2 to search for the optimal helical rise and twist. After CTF refinement and postprocessing, a 3D map at a resolution of 3.8Å was obtained.

To build the 1B^P^ atomic model, one beta rung of the 1B model was fitted into the 1B^P^ density using a rigid-body fit in ChimeraX^51^ (v1.10.1), followed by a jiggle fit and all-atom refinement in Coot^49^ (v0.9.8.96). Position 53 was mutated into a threonine and the model was extended to five beta rungs, which were refined together in phenix.real_space_refine (PHENIX v1.21.2). After refinement, the outer two beta rungs were removed in Coot. NCS and secondary structural restraints were included during the iterative refinement process.

### FLIM of aSyn inclusion pathology

h-FTAA was synthesized and purified as previously described^36^ and paraffin-embedded sections of 1B-injected wild-type mouse brains (129SV) and regions from human donor brain samples were stained with h-FTAA and processed for immunofluorescence against pS129Syn (EP1536Y) for the inclusions or against human aSyn (LB509) for the 1B inoculate and counterstained with the nuclear marker DAPI as previously described^37^. To confirm previous observations regarding the capability of h-FTAA to discriminate Lewy bodies and GCIs using FLIM^37^, human brain samples from one MSA and one DLB subject were obtained from the repository of the University Hospital of Bordeaux (CRB-BBS). Subjects or a legal representative had given informed written consent for collecting and using clinical and post-mortem data, and mandatory regulatory approval for post-mortem brain tissue use was obtained from the French Ministry of Higher Education and Research (MESR AC-2024-6715, DC-2024-6714). FLIM was performed on a Leica SP8 WLL2 confocal microscope on an inverted stand DMI6000 (Leica Microsystems, Mannheim, Germany), using HCX Plan Apo CS2 63X oil NA 1.40 objective. This microscope was equipped with a pulsed white light laser 2 (WLL2) with freely tuneable excitation from 470 to 670 nm (1 nm steps) and a diode laser at 405 nm. A FALCON module enabled us to perform FLIM measurement based on Time Correlated Single Photon Counting (TCSPC) technique. The FLIM analysis was done using phasor plot representation of the data and was focused on pS129Syn-positive inclusions^37^, or LB509-positive regions in the case of the 1B inoculate. For Gallyas silver impregnation, mouse brain sections were processed as previously described^38^ using a commercial Gallyas staining kit (Morphisto, Offenbach am Main, Germany).

### Atomic force microscopy

1B aSyn fibrils were deposited on a freshly cleaved mica disk and left to adhere for 15 minutes in a humid chamber at room temperature. The sample was rinsed and imaged in saline buffer solution (H_2_O, HEPES 20 mM, NaCl 100 mM, pH 7.4). Atomic Force Microscopy imaging (AFM) was performed using the Dimension FastScan setup (Bruker) operating in PeakForce Quantitative Nano-Mechanics (PF-QNM) mode in liquid at room temperature, using sharp SNL-C probes (Bruker, Silicon tips on silicon nitride cantilevers), with a nominal spring constant of 0.24 N/m, a resonance frequency of 56 kHz, and a nominal tip radius of 2 nm. Fibrils were imaged with a peak force frequency of 1 kHz, a scan rate of 0.5 Hz and a constant setpoint force of 500 pN. Image analysis was performed using Nanoscope analysis (Bruker).

### Statistics and Reproducibility

For all experiments/conditions without quantitative analyses, the images shown were representative of at least 3 independent observations. In all other cases, statistical analyses data were presented as means ± SD or otherwise stated in the Figure legends, and were performed using GraphPad Prism (v.9). Numbers of independent samples as well as the statistics tests used are indicated in the figure legends for each experiment. No statistical methods were used to predetermine sample size. Sample size was determined based on our previous publications. Data distribution was assumed to be normal, although this was not tested. Data assignment, organization and collection were randomized. Data collection and analysis were not performed blindly. No animals or datapoints were excluded from the analysis,

## Acknowledgements

Cryo-EM data were in part collected at the Dubochet Center for Imaging (DCI) in Lausanne. The DCI-Lausanne is a joint initiative of the EPFL and the Universities of Lausanne and Geneva. We thank Alex Myasnikov from the DCI-Lausanne for expert technical assistance in cryo-EM data collection, Ænora Letourneur and Fabrizio Cavallaro for their help with primary culture and immunofluorescence, Francesca Serinelli for support with paraffin histology, Sandra Dovero and Nathalie Biendon for support with Pannoramic^TM^ slide scanning, Benjamin Dehay for sharing the AAV-Syn mutants vectors, Marion Picquemal for support with animal surgery, Latifa Lakhdar, Aude de César and Damien Gaillard for animal experiments in M83 mice, Sabrina Lacomme and Fabrice Cordelières from the Bordeaux Imaging Center for their help with sample fixation for CLEM and with the Zeiss CD7 platform respectively. Room-temperature EM was collected in part at the electron microscopy facility (EMF) of the University of Lausanne. We thank Christel Genoud and the EMF team for their assistance. This work was supported by the Swiss National Science Foundation, Grants 310030_188548 & 10.002.679, the Swiss Parkinson Foundation, the Department of Excellence Initiative of the Italian Ministry of Research, the Innovative Medicines Initiative 2 Joint Undertaking under grant agreement no. 116060 (IMPRiND), supported by the European Union’s Horizon 2020 research and innovation program and EFPIA, the French Agence Nationale de la Recherche (grants ANR-21-CE18-0045-03, ANR-22-CE16-0002 and ANR-24-CE93-0003), and the Fondation pour la Recherche Médicale, grant MND202411019711.

## Contributions

D.B. determined 1B structure, performed CLEM experiments and immunogold labeling, wrote manuscript; M.K. performed in vivo experiments in 129SV wild-type mice and neuropathology stainings; L.v.d.H. determined 1BP structure, performed immunogold analysis, wrote manuscript; H.d.LS. produced aSyn and fingerprinted 1B, SEC-purified aSyn, performed the FLIM experiments; A.J.L. performed CLEM experiments; F.D.N. designed experiments, performed in vivo experiments in 129SV wild-type mice and neuropathology stainings; I.M. contributed to cryo-EM structure determination; J.V. performed the analyses of the in vivo experiments in transgenic M83 mice and purified the 1B^P^ fibrils from brains; C.F. performed the AFM of 1B fibrils; M.B. SEC-purified aSyn; ML.A. performed surgery on wild-type mice; A.R. performed post-mortem brain perfusion/fixation of C57BL/6 wild-type mice; E.B. participated in early characterizations of 1B; MH.C. and W.G.M. certified and provided the sections from MSA and DLB patient donors; A.L. SEC-purified aSyn; L.B. participated in early characterizations of 1B; C.P. performed the FLIM experiments; K.P.R.N. synthetized h-FTAA; F.L. contributed to setting up the fibril isolation procedures from mouse brains, participated in early characterizations of 1B; T.B. designed experiments; D.D.L. designed experiments; F.D.G. designed experiments, performed the HCA experiments, as well as the confocal and slide scanning neuropathology analyses of wild-type and M83 mice, and wrote the manuscript; H.S. conceived and managed the project, wrote manuscript; F.I. conceived and managed the project, produced aSyn and fingerprinted 1B, performed the FLIM experiments, performed the HCA experiments, as well as the confocal and slide scanning neuropathology analyses of wild-type and M83 mice, and wrote manuscript. All authors contributed to the paper and discussed the meaning of the results.

## Additional Information

Correspondence and requests for materials should be addressed to Henning Stahlberg and François Ichas.

## Data availability

For the 1B and 1B^P^, the raw cryo-EM images are available at the EMPIAR database, accession numbers EMPIAR-12012 and EMPIAR-12890, respectively. The reconstructed 3D maps are available at the EMDB database, accession numbers EMD-19986 and EMD-54402. The atomic models are available at the PDB database, accession number PDB-9EUU, and PDB-9RZF.

## Extended data figures

**Extended Data Fig. 1.**
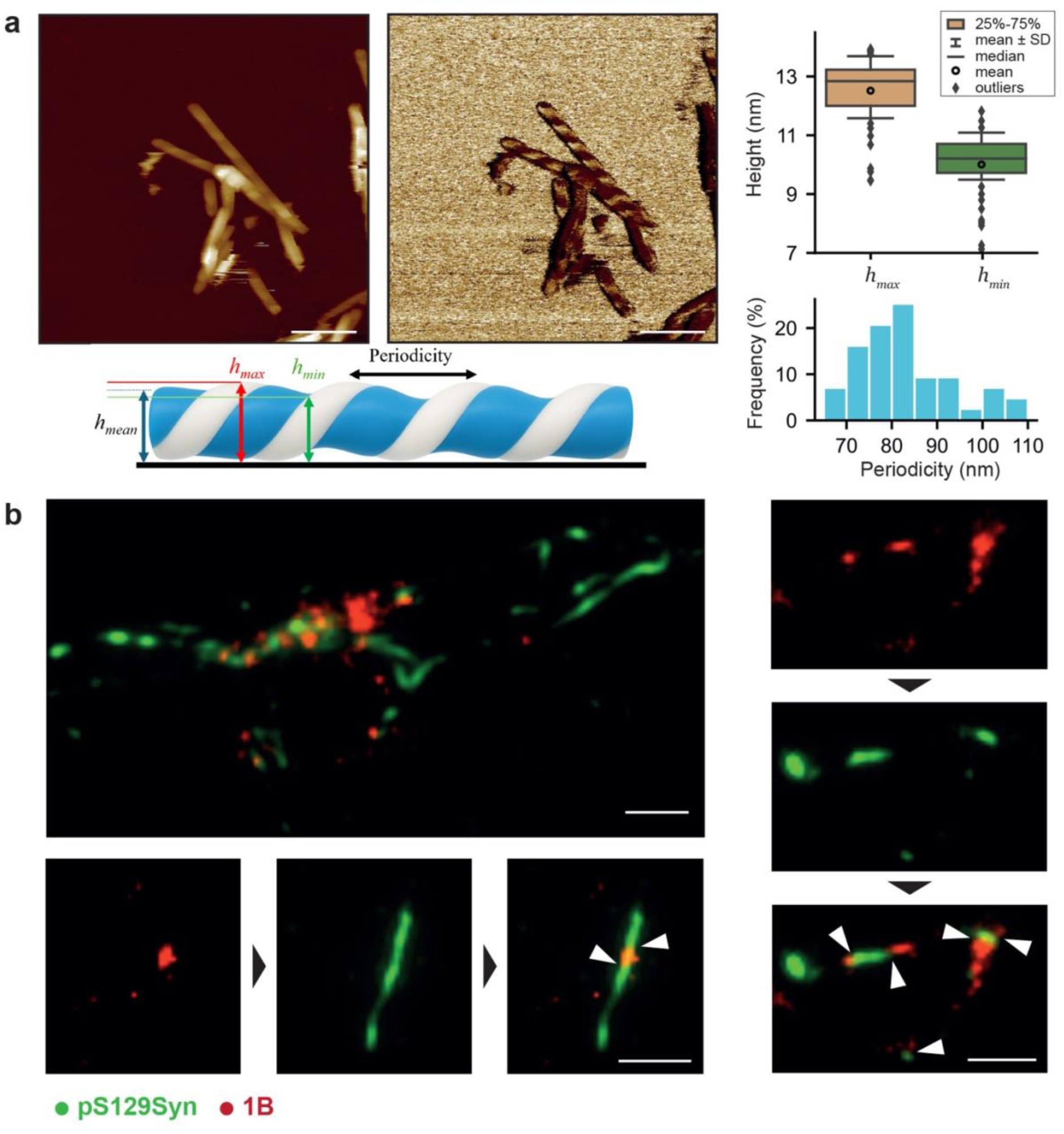
AFM appearance of 1B fibrils and early formation of “seeding junctions” between 1B and the nascent seeded inclusions in neuronal processes. **a.** Topographical (Height Sensor) and nano-mechanical mapping (Derjaguin-Müller-Toporov (DMT) Modulus, log representation) of 1B fibrils using atomic force microscopy (AFM) in liquid. Right panels: fibril height values (see cartoon for definitions) for n = 103 measurements (plot population statistics descriptors as shown in plot inset), and left twist periodicity for n = 12 fibrils. Scale bars = 240 nm. **b.** Examples of early “seeding junctions” between 1B and the nascent seeded inclusions in neuronal processes two weeks post seeding (arrowheads = seeding junctions). 96-well pooled primary cultures of cortical neurons from 8 wild-type mouse embryos (C57BL/6) were challenged at day-in-vitro 7 (DIV7) with 10 nM 1B fibrils (equivalent aSyn monomer concentration). The neurons were fixed two weeks later at DIV24 and both the 1B input fibril remnants (red) and the seeded aSyn inclusions (green) were revealed. Indeed, we took advantage of the fact that synthetic 1B is made of human aSyn and thus that it can be specifically recognized by LB509 which is a human aSyn-specific antibody. Mouse aSyn is LB509 negative. The seeded inclusions were revealed contemporary using EP1536Y against pS129 aSyn. This allowed us to observe many seeding junctions in the neuronal processes (arrows). Note that in a majority of images, and in agreement with early reports, the seed (1B in our case) is not phosphorylated, and that the growing seeded inclusions are instead pS129Syn-positive. At later time points 1B seeds get overwhelmed and are no longer discernible in the mass of the seeded inclusions (not shown). The disappearance of the 1B seeds is even faster *in vivo* (within a few days, not shown). (scale bars = 10 µm)

**Extended Data Fig. 2.**
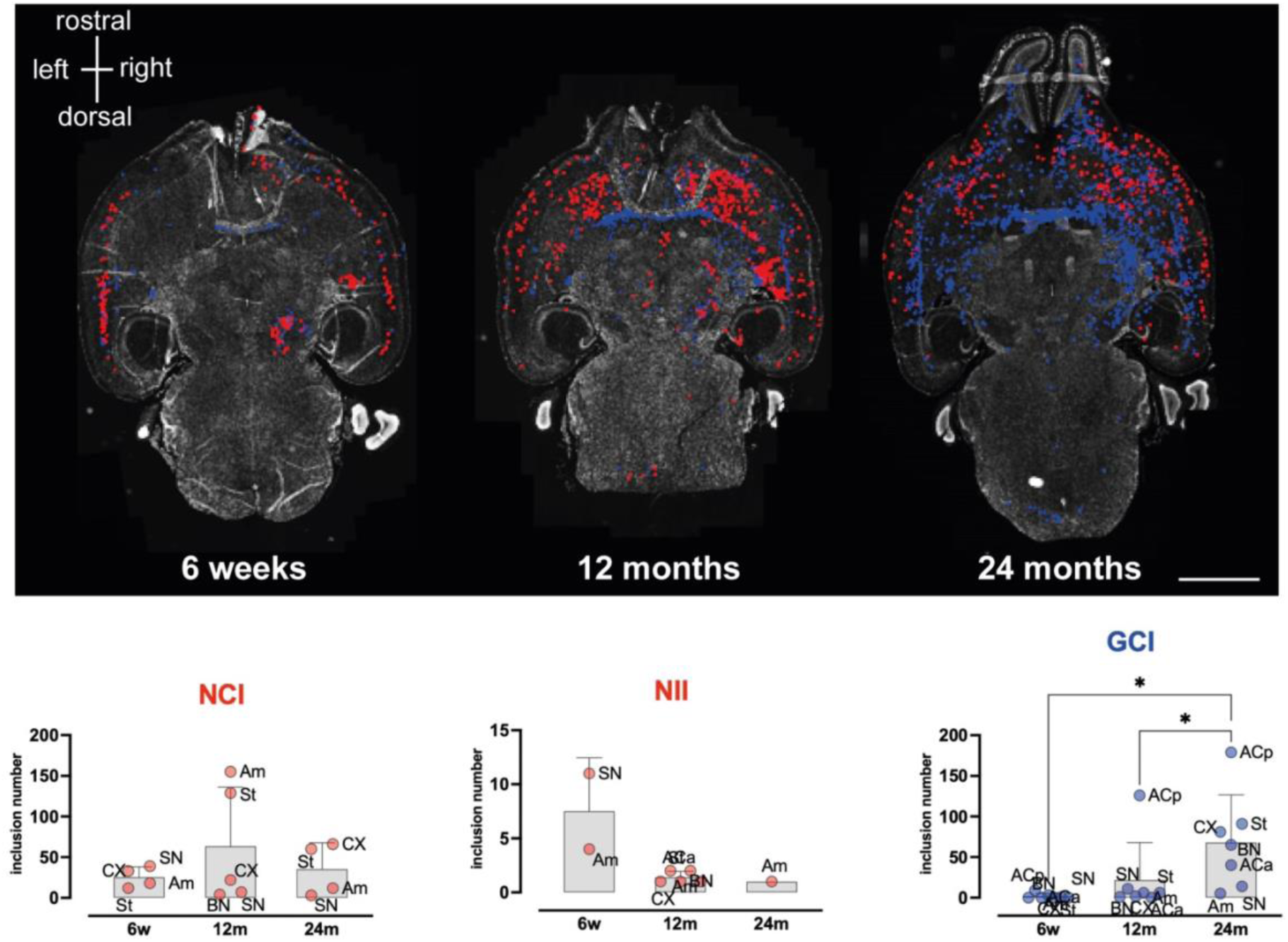
Somatic inclusions seeded by 1B *in vivo* observed at 3 time points. Depending on the neuroanatomical structure and the time point under consideration, experimental GCIs (blue) coexisted with a variable load of surrounding neuronal somatic aSyn inclusions (NCIs & NIIs, both in red), which represent the forefront of the somatic aSyn pathology at earlier time points to progressively make way to a growing population of GCIs. The brain sections are the ones shown in **Fig. 1f** and were stained with DAPI for nuclei, Sox10 to identify the oligodendrocytes, and EP1536Y for pS129Syn. All the inclusions were manually annotated according to these 3 markers. Anatomical quantification regions were as follows: Am: amygdala; ACa: anterior limb of the anterior commissure; ACp: posterior limb of the anterior commissure; BN: bed nucleus of the stria terminalis; CX: cortex; St: caudate putamen (striatum); SN: *substantia nigra*. In agreement with visual perception, the neuronal inclusions peak at 12 months in the Caudate putamen (St) and the Amygdala (AM) to decrease at 24 months, while the GCIs suddenly increase in the posterior limb of the Anterior Commissure at 12 months, and then in most of the other brain regions. Note that NIIs form especially early (at 6 weeks) in the SN and the AM, then in other regions at 12 months, to eventually disappear but in the AM at 24 months. The mean number of GCIs in all regions at 24 months is significantly increased compared to the 2 other time points (Mean ± SD of the regions shown together with the individual region inclusion number values, * = one-way ANOVA p<0.05: p = 0.0462 for 6w/24m, p = 0.0278 for 12m/24m)

**Extended Data Fig. 3.**
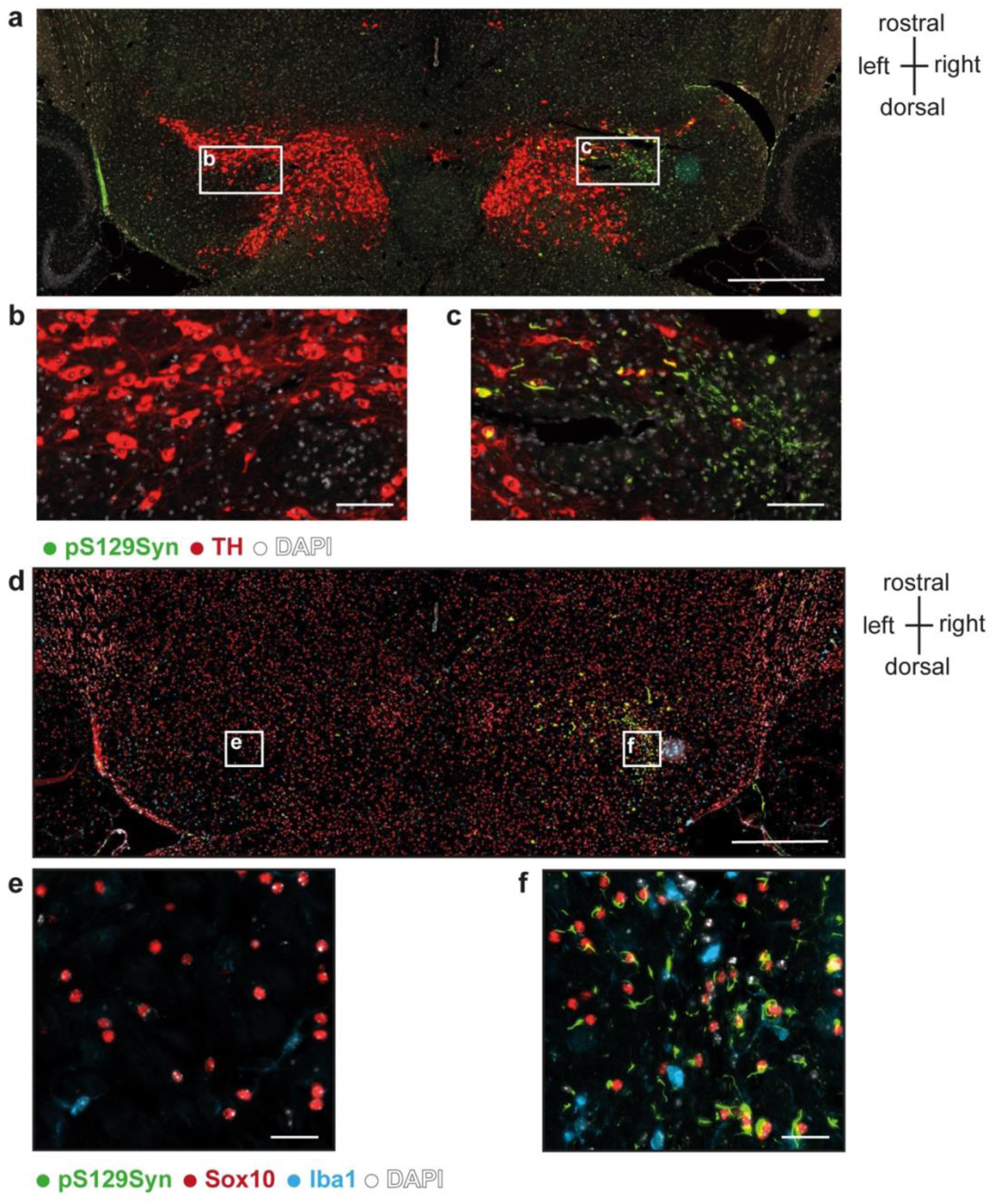
Semi-topographic view of a horizontal brain section from a C57BL/6 wild-type mouse injected 17 months earlier at the level of its right substantia nigra (SN) with 1B fibrils. **a.** aSyn inclusions are revealed using the anti-phospho aSyn antibody #64 (green), dopaminergic neurons using an anti-tyrosine hydroxylase antibody (TH, red) and nuclei using DAPI (white). Left and right SNs are visible with a loss of immunoreactivity symmetry with depletion of TH-positive neurons and unilateral aSyn inclusions on the right side. The TH-depleted/aSyn inclusion-rich region is enlarged in b, the corresponding left region is enlarged in c. d. Semi-topographic view of the n+1 horizontal section consecutive to (a). aSyn inclusions in green (anti-phospho aSyn antibody #64), OLs in red (anti-Sox10 antibody), microglial cells in blue (anti -Iba1) and nuclei in white (DAPI). The right SN enlarged in f contains numerous GCIs and sparse microglial cells. e. f. Enlarged view of the corresponding region of the left SN. Scale bars: a, d = 500 µm; b, c = 100µm; e, f = 20 µm.

**Extended Data Fig. 4.**
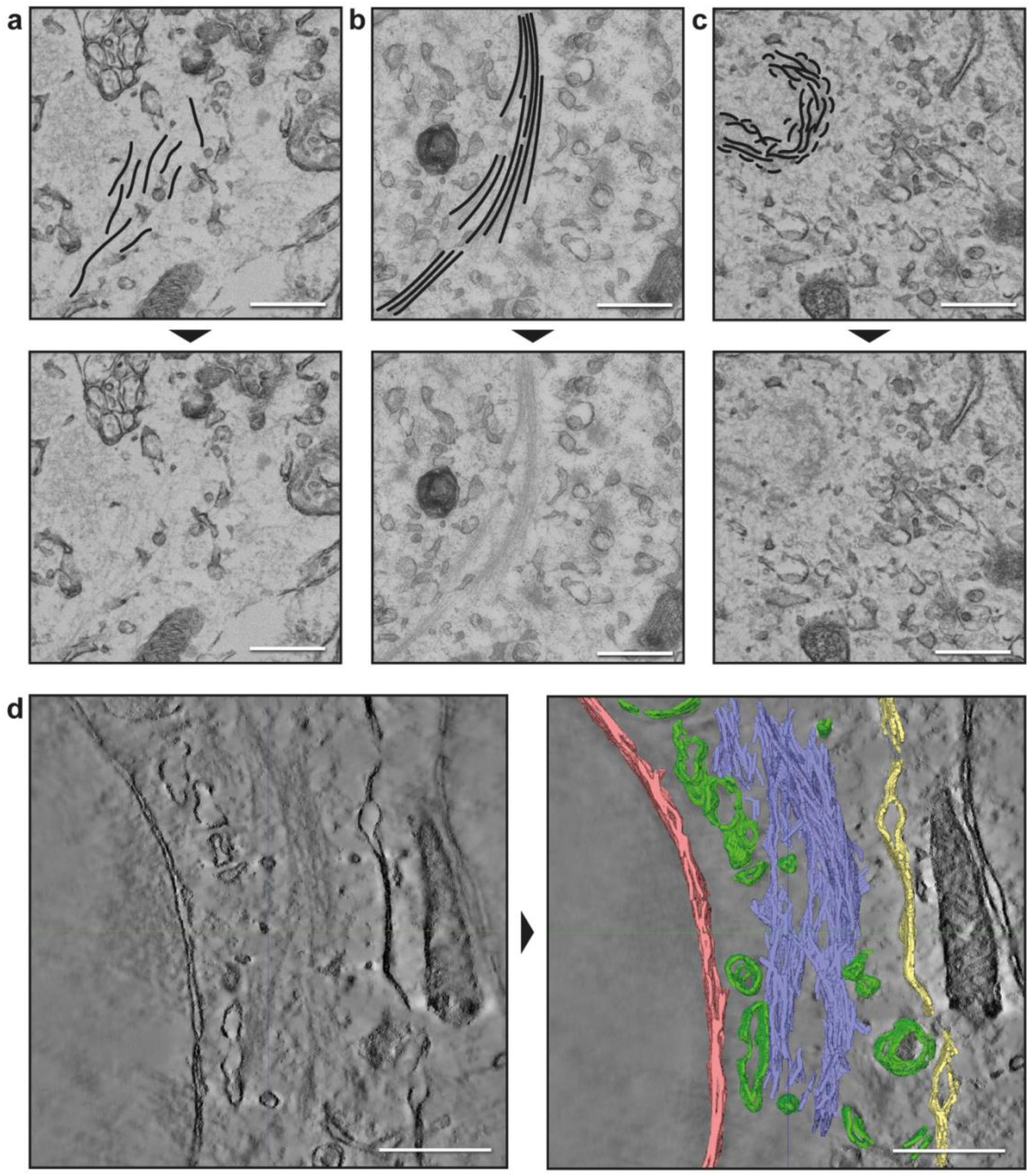
Tissue ultrastructure and tomography of 1B-seeded mouse brain. **a-c.** High magnification EM images of different types of fibril arrangements, indicated by hand made drawings in the upper row panels, and without indication in the lower row panels. **a:** “dispersed”, **b:** “bundled”, and **c:** “clustered”. **d.** Tomogram segmentation of aSyn fibril bundles. Tomograms were manually segmented to visualize the fibril bundling (blue), the nuclear membrane (red), the cell membrane (yellow), and other membranous compartments (green). Top: A slice view of the reconstructed tomogram with applied filtering to better visualize the filaments within the inclusion. Bottom: Overlay with 3D-segmented elements. Scale bars: 500 nm.

**Extended Data Fig. 5.**
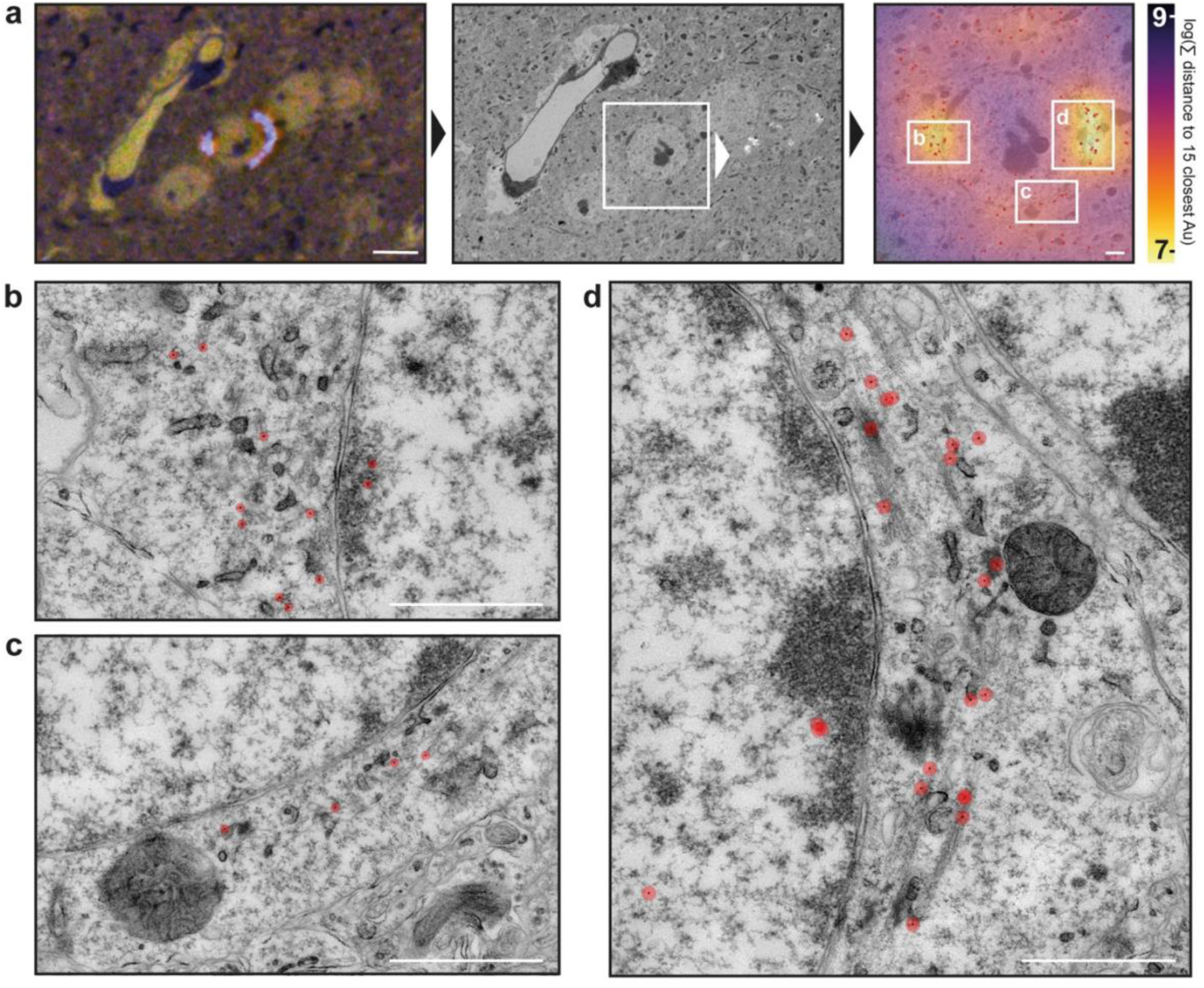
Immunogold analysis against aSyn in the mouse brain. **a.** Immunohistochemistry (IHC) labelling (inset) indicates two regions with aSyn pathology wrapping around the nucleus of an affected cell. The correlated EM image with on-grid immunolabelling against aSyn (left) shows an increase of gold particles in comparison to the background as indicated by the heatmap (right). Higher magnifications are indicated by the labelled boxes. **b.** Zoomed view on the left side of the nucleus, highlighted are the gold particles in red, laying on top of dispersed fibrils. **c.** Lower part of the pathology from the right side of the cell. The zoomed area shows four gold particles laying on top of fibril pathology. **d.** Zoom of right-side pathology, with highest concentration of gold particles, laying on fibril bundles. Scale bars: in a = 5 µm (left) and 1 µm (right); in b-d: 1 µm.

**Extended Data Fig. 6.**
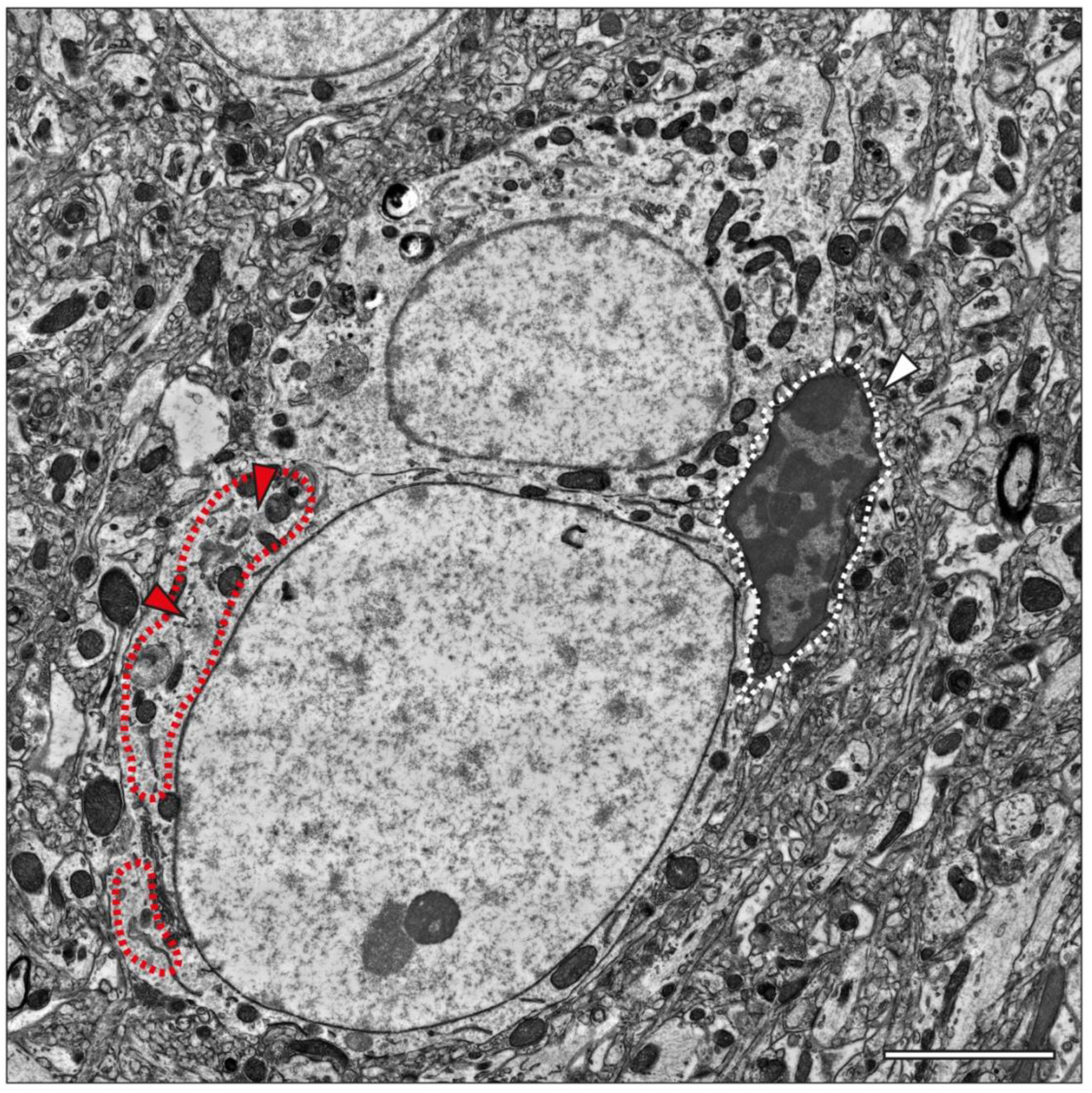
Pathological oligodendrocyte in direct contact with other cells. Cells with a distinct morphology, previously described as “dark microglia”, can be found in close proximity or in direct contact with pathological cells (white arrowhead). The total cell volume of the dark microglia is indicated by the white dashed line. In this specific case the fibrillar pathology of the oligodendrocyte appears in a dispersed arrangement with two smaller clusters (red arrowheads). Total area of immunopositive IHC is indicated by a red dashed line. Both oligodendrocytes contain aSyn pathology however for the upper cell volume it is not visible in this EM slice (based on IHC positive signal from other EM grids). Scale bar = 3 μm.

**Extended Data Fig. 7.**
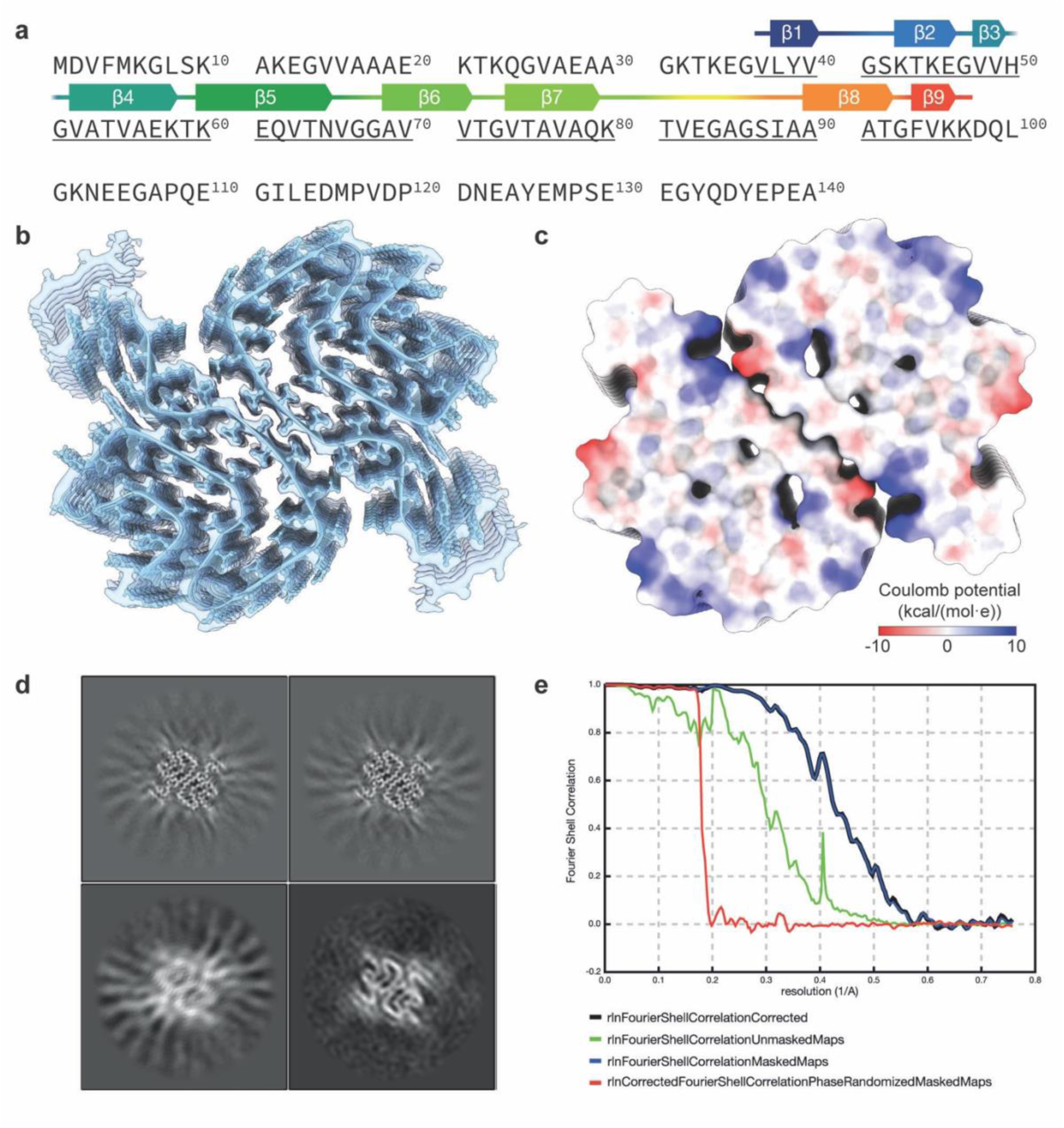
Further structure analysis of the 1B conformation. **a.** Sequence of aSyn with the resolved density indicated by the colored line. Secondary beta-sheets are indicated in consecutive order by β1-9. **b.** The final 1.94 Å map of 1B with the model fitted. The extra densities flanking the starting stretches were not built. **c.** Surface color representation by the electrostatic potential. **d.** 3D class averages from early processing stages indicate a homogeneous sample. **e.** FSC curve of 1B with cutoff at 0.143 of the final map at 1.94 Å.

**Extended Data Fig. 8.**
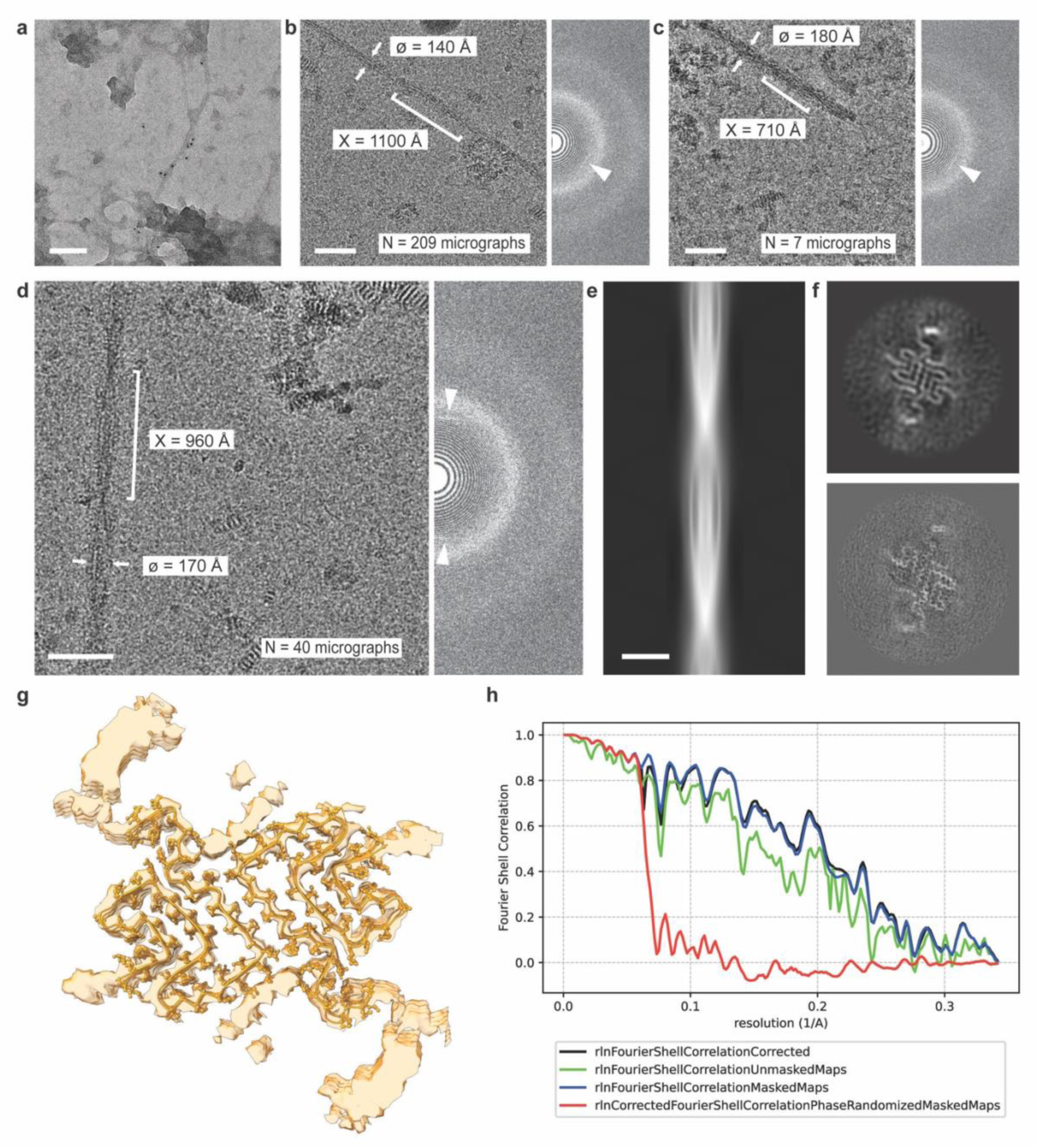
Cryo-EM analysis of 1B^P^ fibrils purified from a tg M83+/-mouse brain seeded with 1B fibrils. **(a)** Negative staining combined with immunolabeling against aSyn-pS129 reveals that, amid aggregated precipitates, individual aSyn fibrils are present. **(b-d)** Cryo-EM micrographs (left) of intact fibrils observed in the sarkosyl-insoluble fraction, together with their power spectral density (right). A reflection at ∼4.8Å (white arrowheads) reveals that these are amyloid fibrils. Presumably, the fibrils in (b), (c) and (d) represent 3 distinct conformations as can be seen from their different morphologies, although the particle counts are too low to verify this with a 3D reconstruction. The crossover distance (X) and largest diameter (ø) are indicated, as well as the number of micrographs (N) displaying at least one crossover of a fibril of the same morphological group as determined by manual inspection. The fibrils in group (d) did eventually yield a 3.8Å structure, while those in groups (b) and (c) did not yield any structure despite many attempts. **(e)** Reprojection of the 2D inimodel constructed from 2’492 particles selected from group (d). **(f)** 3D helical reconstruction of 1’913 particles selected from group (d), before (top) and after (bottom) post-processing in Relion. **(g)** The final 3.8Å map of 1B^P^ with the model fitted. **(h)** FSC curve of 1B^P^. Scalebars: (a) 100 nm, (b-d) 50 nm, (e) 20 nm.

**Extended Data Fig. 9.**
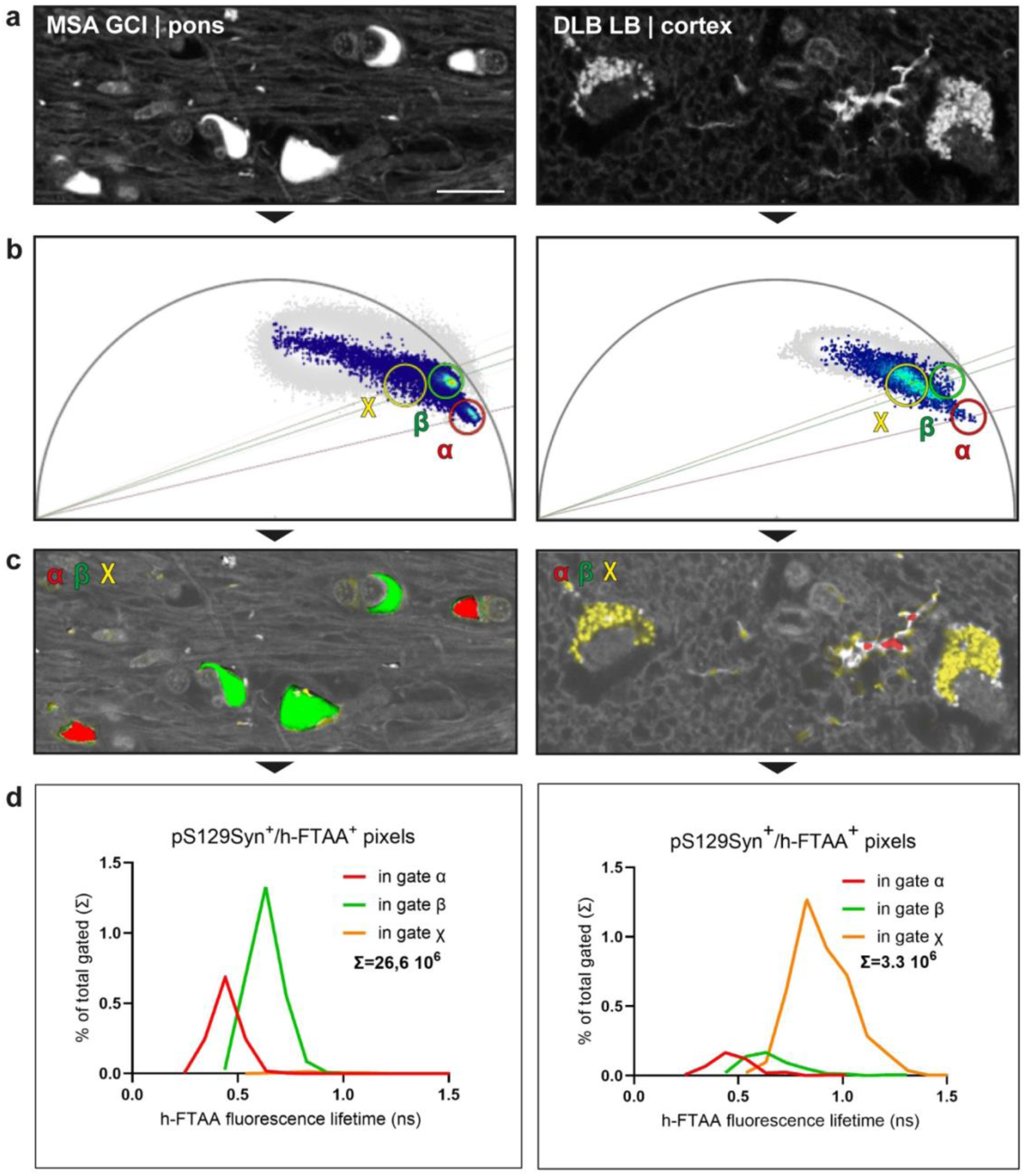
h-FTAA is a conformation-sensitive probe that enables discrimination between aSyn fibril strains associated with MSA and DLB through FLIM of brain tissue sections. It was previously reported that FLIM of h-FTAA can be used in histological brain sections to differentiate the aSyn fibrils populating Lewy bodies (LBs) from those populating GCIs in MSA^37^. These findings established h-FTAA as a conformation-sensitive amyloid probe capable of discriminating distinct aSyn fibril strains in human brain sections via FLIM^37^. In agreement with these previous observations, FLIM of h-FTAA allowed us to derive fibril strain-specific phasor plot signatures directly on brain tissue sections. Using this approach, we confirmed that h-FTAA FLIM could indeed discriminate the pS129Syn-positive inclusion pathology of MSA vs. DLB (a: h-FTAA intensity images - pS129Syn not shown) and used the disease-specific hotspots in the respective FLIM phasor plots to define 2D quantification gates (b,c) (ɑ, β: MSA-specific; χ: DLB-specific). The fluorescence lifetime of the gated pixels is shown in d for respectively GCIs (26,6 million inclusion pixels gated) and LBs (3.3 million inclusion pixels gated). In agreement with previously published data^37^, the distribution histograms make it clear that the majority of pixels belonging to LBs are characterized by a fluorescence lifetime which is in the nanosecond range, while those populating GCIs show an approximately halved fluorescence lifetime with 2 distinguishable populations. Scale bar = 20 μm.

**Extended Data Fig. 10.**
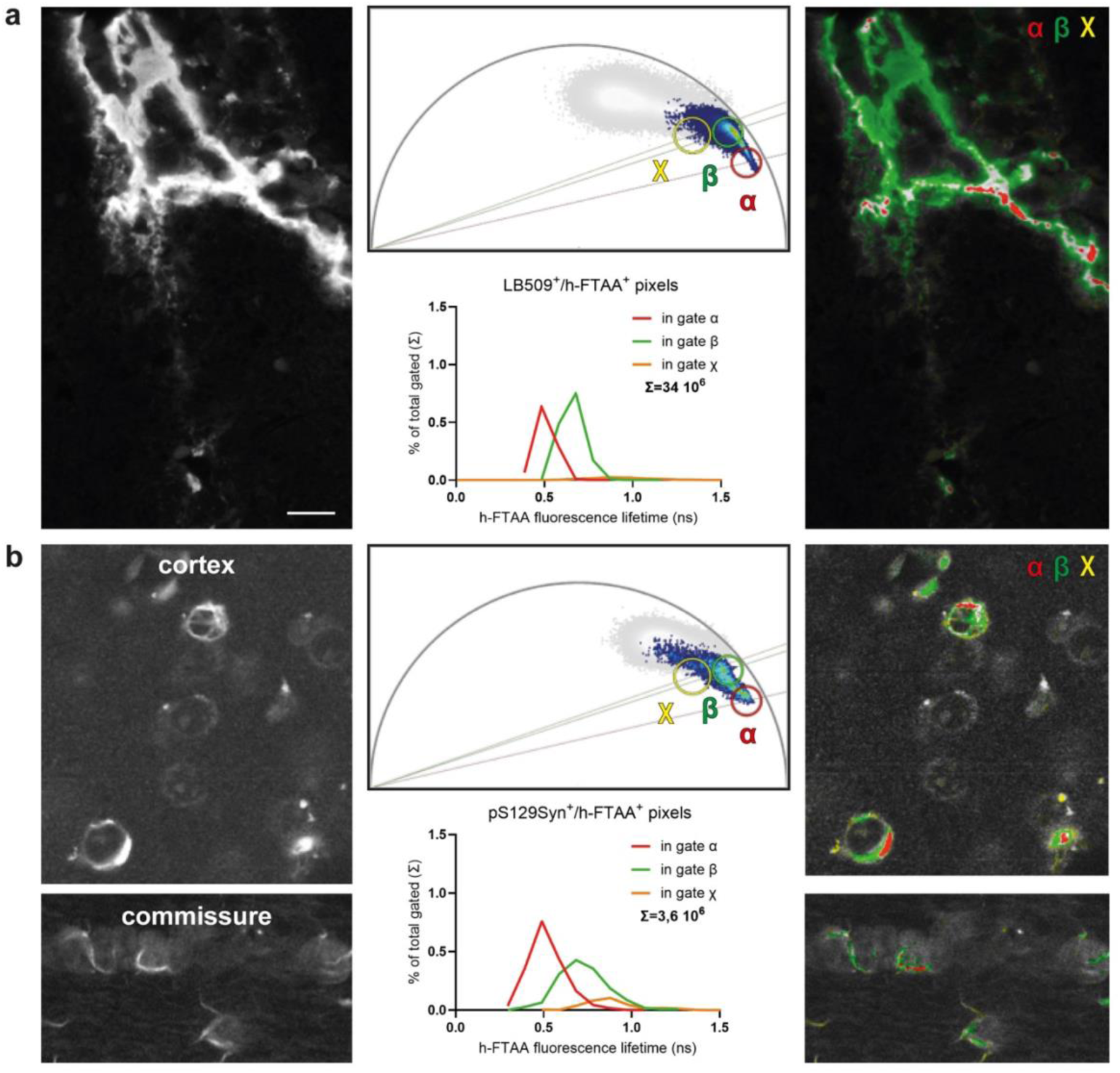
*In situ* h-FTAA FLIM indicates that both the input 1B fibrils and the resulting inclusions seeded in wild-type mice exhibit a high degree of conformational similarity with the MSA strain. The phasor plot gates discriminating GCIs and LBs described in Extended Data Fig. 9 were applied in the phasor plots of the intracerebral 1B fibril inoculate (positive to the anti-human aSyn antibody LB509, not shown), still observable in the CP of wild-type mice 24 hours after injection (**a,** from left to right: h-FTAA intensity image, corresponding phasor plot with gates and quantifications, distribution of gated pixels in image) or in the plots characterizing the inclusions that formed in neurons and OLs during the 16 months following 1B injection (pS129Syn-positive, not shown) (**b**). The number of inclusion pixels quantified by FLIM and gated in ɑ, β, and χ was >10^6^ in each case, and the distribution of the pixel population in the phasor gates ɑ+β, and χ was eventually used to score the similarity of the different fibril inclusion types (Supplementary Information Table 3). Scale bar = 20 μm.

**Extended Data Fig.11.**
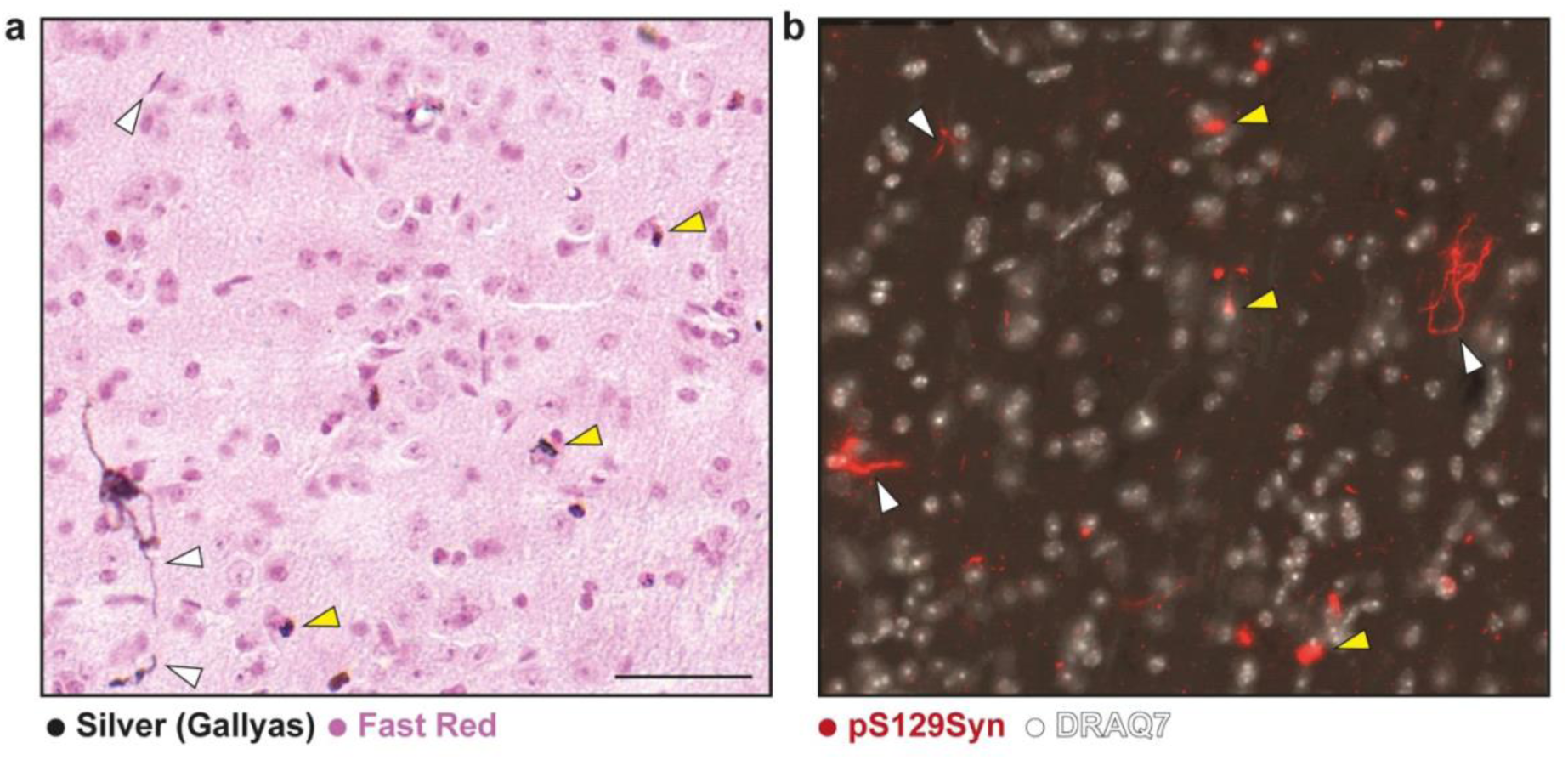
The pathology seeded by 1B is Gallyas-positive. The MSA-specific silver impregnation technique of Gallyas highlights somatic (yellow arrowheads) and neuritic (white arrowheads) inclusions which developed during 16 months in the substantia nigra of WT mice injected with 1B (**a**); other section in the same region reveals pS129Syn-positive inclusion pathology (**b**). Scale bar = 100 μm.

**Extended Data Fig. 12.**
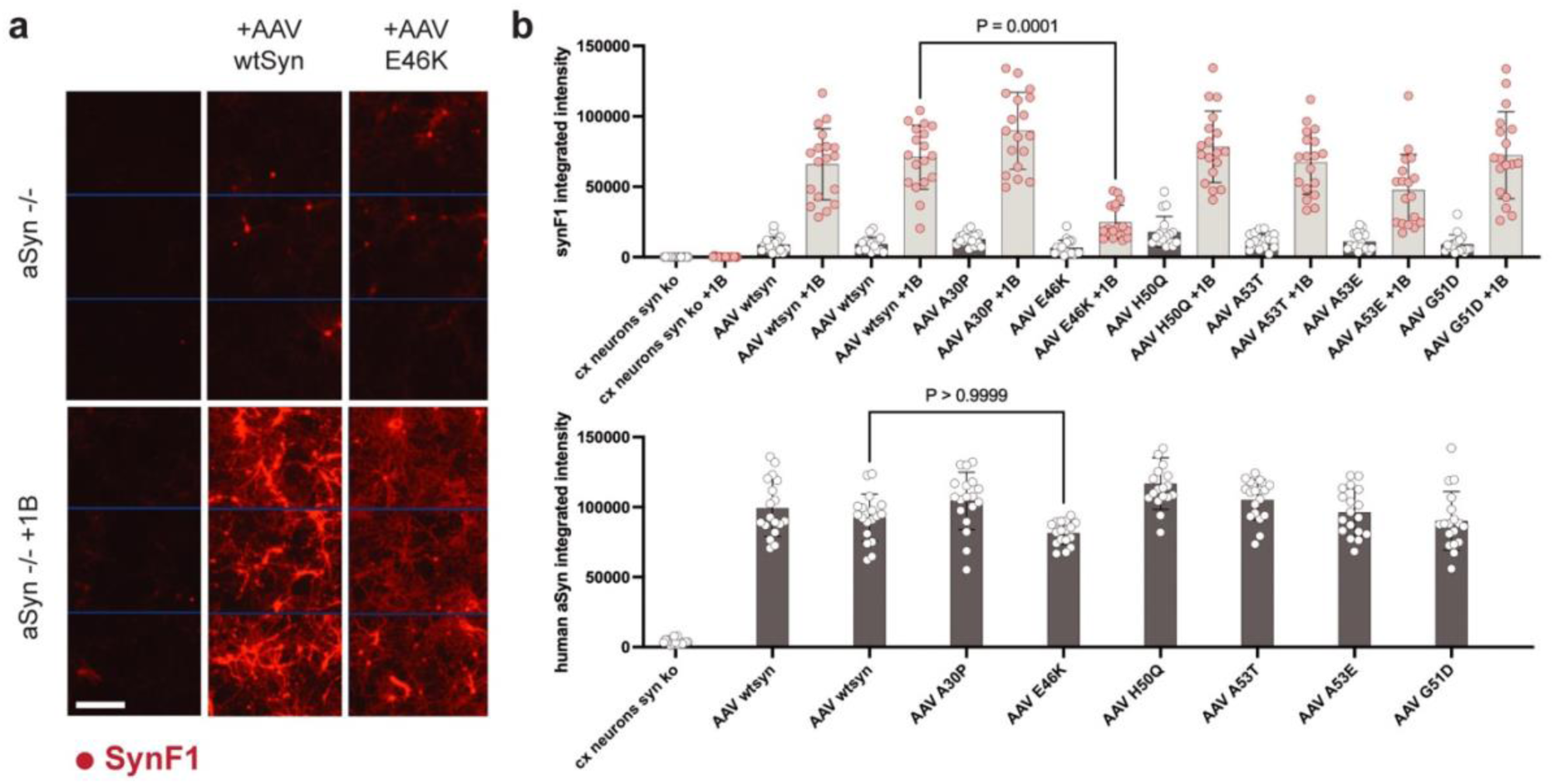
Inclusion seeding by 1B is inhibited by E46K which disrupts the pseudo-Greek-key motif stem stabilizer (E46-K80 salt bridge), but not by aSyn mutations in the inter-protofilament interface. **a.** 96-well pooled primary cultures of cortical neurons from 8 aSyn-KO mouse embryos were challenged or not at day-in-vitro 7 (DIV7) with 10 nM 1B fibrils (equivalent aSyn monomer concentration), and infected or not with AAV-aSyn or AAV-E46K aSyn at DIV11. The neurons were fixed at DIV30 and the aSyn inclusions revealed using the conformation-dependent antibody SynF1 (red); three immunofluorescence fields of views (FOVs) imaged in a test well for each condition, illustrate: (i) the absence of SynF1and human aSyn signal at DIV30 in the absence of endogenous aSyn (aSyn-/-line, left column) , (ii) aSyn inclusion seeding by 1B in conditions of aSyn expression (aSyn -/-+ 1B line, middle and right column), and (ii) a minor inclusion seeding when E46K aSyn is expressed instead of wild-type aSyn (aSyn -/-+ 1B line, right versus middle column). **b.** High Content Analysis of the experiment shown in (a) and extended to a series of aSyn mutants based on 18 FOVs from 6 wells for each condition. Mean ± SD as well as individual values are plotted for each condition The AAV-aSyn condition was doubled to document robustness. Kruskal-Wallis test followed by a post-hoc Dunn’s test for multiple comparisons indicates that E46K aSyn is the only mutation reducing inclusion seeding by 1B (p= 0.0001). In the AAV-only conditions, human aSyn deriving from AAV infection was quantified using the human aSyn-specific antibody MJFR1 (no endogenous mouse aSyn is present in these neurons). The results were compared with the same test and post-hoc test combination as before, which indicated the absence of statistically significant differences in terms of levels of expression driven by the different AAVs. In particular, no statistically significant difference existed between AAV wt aSyn and AAV E46K aSyn in terms of expression levels, indicating that the difference seen with SynF1 after seeding with 1B was not due to a lesser expression of E46K aSyn. Scale bar = 100 μm.

**Extended Data Fig. 13.**
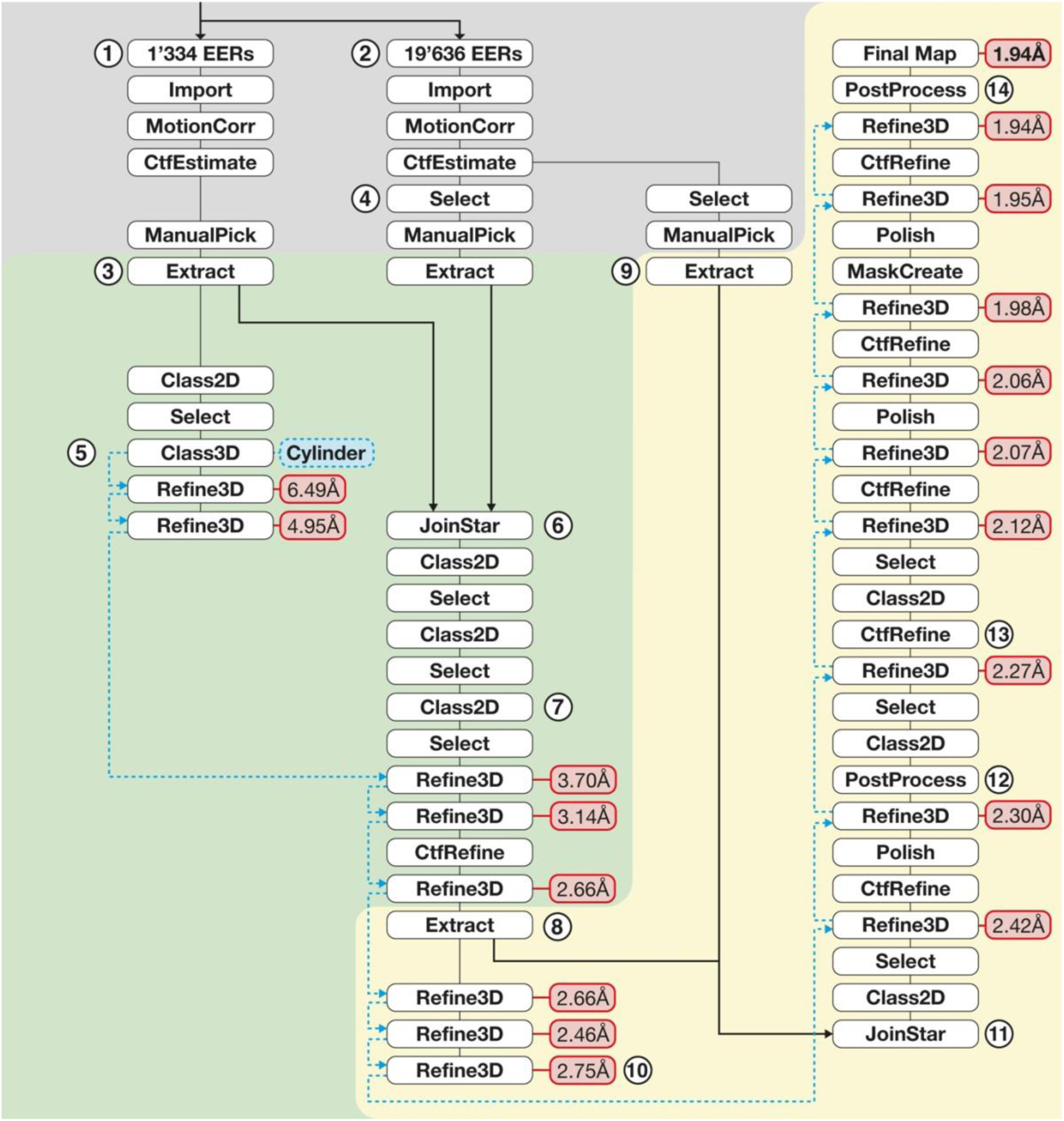
Processing overview (schematic) for 1B reconstruction. Detailed processing overview representing individual jobs within RELION (white boxes), with the black line indicating subsequent steps. Red boxes indicate estimated resolution at given Refinement steps. Blue dotted line represents the reference volumes used. The background color represents data preprocessing (grey), binned processing (green), and unbinned processing (yellow). **1.** During data acquisition the first 1’334 micrographs (Electron Event Recordings) were processed during the session, to assess the quality of the dataset on the fly. **2.** The remaining 19’636 micrographs were processed after the data acquisition had completed and were later introduced to the dataset at step 6 and 11 respectively. **3.** For initial analysis manually traced fibril segments were extracted with a box size of 600px, binned to 300px. **4.** From the full dataset selections were made based on the estimated CTF resolution to reduce the amount of manual screening of the full dataset. As such two groups were analyzed: Add values. Manual picking of particles was done in parallel to the processing of the extracted particles from 3. **5.** Initial 3D-Classification was done with a featureless cylinder as a low-pass filtered reference volume. **6.** The first set of additional particles from the full data set was introduced to the processing. **7.** The full particle batch was cleaned by multiple rounds of 2D classification and was subsequently refined to 2.66Å with a reference volume from the previously obtained 4.95Å structure from the initial screening. **8.** With a well-aligned structure, the data was transitioned to unbinned processing, and the respective particles were re-extracted. **9.** The second set of additional particles from the full dataset was extracted at the unbinned box size. No further manual selection of the full dataset was made. **10.** After transitioning to the unbinned box size the refinement was redone on the 2.66Å structure. **11.** All particles are joined together for the final processing. **12.** The structure was subjected to Refinement and PostProcessing with intermediate steps of Polish and **13.** CTF refinement until the final high-resolution structure was obtained. **14.** Processing concluded when no further improvement in the iterative process could be made. The model was built into the final map at 1.94Å resolution.

## Videos: Confocal animations/3D reconstructions

**SI-Video-1_Fig1a_to_e_GCIs_brain_section_hovering-ACPL-BNST-GCIs:**

Section hovering overview corresponding to **Fig. 1 panels a-e**: DAPI in blue, pS129Syn in white, Sox10 in pink, Iba1 in yellow (only in AC)

**SI-Video-2_Fig5b_upper_right_NII_and_NCI_coexisting_in_dopam_neuron-LeSTRIAP7-99c:**

3D animation of NII & NCI shown in **Fig. 5 panel b, upper right**: DAPI in white, pS129Syn in green, Tyrosine Hydroxylase in red

**SI-Video-3_Fig5b_upper_left_NII_in_a_dopamiergic_neuron-LeSTRIAP7-85:**

3D animation of NII shown in **Fig. 5 panel b, upper left**: DAPI in white, pS129Syn in green, Tyrosine Hydroxylase in red

**SI-Video-4_NCI_and_NII_in_dopaminergic_neuron_without_nuclear_marker-LeSTRIAP7-97:**

3D animation of a Tyrosine Hydroxylase (red) positive cell from the s*ubstantia nigra* of the mouse sampled in **Fig. 5**. pS129Syn is in green. Both NCIs and NIIs can be observed without need for a nuclear marker

**SI-Video-5_Fig5b_lower_left_Neuron_to_glia_protrusion_of_NCI-LeSTRIAP7-99a:**

3D animation of **Fig. 5 panel b, lower left**: The NCI of the cortical neuron (bigger nucleus) seems to protrude within the interior of the lining glial cell (smaller nucleus): DAPI in white, pS129Syn in green, Tyrosine Hydroxylase in red (not expressed in this region)

**SI-Video-6_Fig5b_lower_left_Neuron_to_glia_protrusion_of_NCI-LeSTRIAP7-99b**

“in silico” section of 3D animation of **Fig. 5 panel b, lower left**: The NCI is in continuity with a glial nuclear inclusion (GNI): DAPI in white, pS129Syn in green, Tyrosine Hydroxylase in red (not expressed in this region)

## Supplemental Tables

**Supplemental Table 1.** Structural comparison of different aSyn folds with 1B. Structure comparison on the basis of the r.m.s.d. value, as shown in Fig. 3. The fully modelled structures were aligned to the 1B conformation with the matchmaker function in ChimeraX. The best fit subunits were then further compared on three regions on the Cα r.m.s.d. values in respect to the different pocket regions and intermediate sequence based on ChimeraX r.m.s.d. measurements. A r.m.s.d. of less than 1 Å suggests high similarity in the structural conformation.

| Comparing 1B (PDB: 9EUU) with: | PDB Entry | Total Pairs Nr. | Total RMSD of one protofilament [Å] | N-Pocket RMSD (res. 39-47), [Å] | Intermediate RMSD (res. 48-87), [Å] | C-Pocket RMSD (res. 88-94), [Å] |
| --- | --- | --- | --- | --- | --- | --- |
| H50Q Mutation | 6PEO | 61 | 0.566 | 0.504 | 0.547 | 0.693 |
| MSA seeded | 7NCK | 61 | 0.781 | 0.597 | 0.815 | 0.892 |
| 1B<sup>P</sup> | 9RZF | 61 | 1.433 | 0.582 | 1.642 | 0.357 |
| MSA purified | 6XYQ | 61 | 2.425 | 3.685 | 1.228 | 0.398 |
| 1-121 aSyn | 6H6B | 58 | 4.856 | 3.418 | 3.484 | 9.225 |
| wt aSyn | 80QI | 56 | 5.282 | 7.631 | 3.171 | 9.532 |
| ThT bound | 7YNM | 61 | 4.688 | 4.643 | 3.225 | 8.511 |

**Supplemental Table 2.** Cryo-EM data collection, refinement, and validation statistics

|  | **1B fibril** | **1B<sup>P</sup> fibril** |
| --- | --- | --- |
| Raw images | EMPIAR-12012 | EMPIAR-12890 |
| Reconstructed volume | EMD-19986 | EMD-54402 |
| Atomic model: | PDB-9EUU | PDB-9RZF |
| **Data collection and processing** |  |  |
| Nominal Magnification | 120'000x | 165'000x |
| Voltage (kV) | 300 | 300 |
| Total electron dose (e/Å<sup>2</sup>) | 40 | 50 |
| Defocus range (μm) | -0.8 to -1.8 | -0.8 to -2.2 |
| Pixel size (Å) | 0.66 | 0.732 |
| Symmetry imposed | C1 with 2<sub>1</sub> cork-screw symmetry | C1 with 2<sub>1</sub> cork-screw symmetry |
| Micrographs | 11'879 | 42'812 |
| Initial particle images (no.) | 108'543 | 3'556 |
| Final particle images (no.) | 51'272 | 1'913 |
| Map resolution (Å)<br>FSC threshold at 0.143 | 1.94 | 3.8 |
| Map resolution range (Å) | 1.88-2.56 | 3.45-31.38 |
| Estimated map sharpening B-factor (Å<sup>2</sup>) | -21.06 | -2.52 |
| Helical twist (°) | 179.52 | 179.55 |
| Helical rise (Å) | 2.43 | 2.4 |
| **Refinement** |  |  |
| Model resolution range (Å) | 1.7 - 2.4 | 3.3 - 9.0 |
| Model composition |  |  |
| Non-hydrogen atoms | 3400 | 2538 |
| Protein residues | 496 | 366 |
| Ligands | 0 | 0 |
| B factor (Å<sup>2</sup>)<br>Protein | 21.06 | 56.98 |
| R.m.s. deviations |  |  |
| Bond lengths (Å) | 0.001 | 0.004 |
| Bond angles (°) | 0.49 | 0.858 |
| Validation |  |  |
| MolProbity score | 1.52 | 1.83 |
| Clashscore | 5.57 | 7.84 |
| Poor rotamers (%) | 0 | 0 |
| Ramachandran plot |  |  |
| Favored (%) | 96.61 | 94.07 |
| Allowed (%) | 3.39 | 5.93 |
| Disallowed (%) | 0 | 0 |

**Supplemental Table 3.** In situ comparison of 1B inoculate, 1B-seeded inclusions, GCIs in MSA and LBs in DLB using h-FTAA FLIM of human and wild-type mouse brain tissue sections. Inclusion pixels quantified by FLIM and gated in ɑ, β, and χ in **Extended data Fig. 10** (>10^6^ in each case) were used for comparison of inoculate and inclusions. For each condition, the % of the inclusion pixel population situated in the phasor gates ɑ+β (MSA specific), and χ (DLB specific) (see **Extended data Fig. 9**), was used to score the similarity of the different fibril inclusions types using the Jaccard similarity index^52^. Similarity matrix analysis demonstrates that both the input 1B fibrils and the resulting inclusions seeded in wild-type mice exhibit a high degree of conformational similarity which is shared with the MSA fibrils, and distinct from the DLB ones. Color coding: green = similar, yellow = dissimilar.

|  | **MSA**<br>GClS<br>pS129Syn+ | **1B-seeded**<br>NCIs & GClS<br>pS129Syn+ | **1B inoculate**<br>LB509+ | **DLB**<br>LBs<br>pS129Syn+ |
| --- | --- | --- | --- | --- |
| **MSA**<br>GClS<br>pS129Syn+ | 1.000 | 0.900 | 0.762 | 0.150 |
| **1B-seeded**<br>NCIs & GClS<br>pS129Syn+ | 0.900 | 1.000 | 0.848 | 0.197 |
| **1B inoculate**<br>LB509+ | 0.762 | 0.848 | 1.000 | 0.259 |
| **DLB**<br>LBs<br>pS129Syn+ | 0.160 | 0.197 | 0.259 | 1.000 |

